# Long-Distance Trail Running Induces Inflammatory-Associated Protein, Lipid, and Purine Oxidation in Red Blood Cells

**DOI:** 10.1101/2025.04.09.648006

**Authors:** Travis Nemkov, Emeric Stauffer, Francesca Cendali, Daniel Stephenson, Elie Nader, Mélanie Robert, Sarah Skinner, Monika Dzieciatkowska, Kirk C. Hansen, Paul Robach, Guillaume Y Millet, Philippe Connes, Angelo D’Alessandro

## Abstract

Ultra-endurance exercise places extreme physiological demands on oxygen transport, yet its impact on red blood cells (RBCs) remains underexplored. We conducted a multi-omics analysis of plasma and RBCs from endurance athletes before and after a 40-km trail race (MCC) and a 171-km ultramarathon (UTMB^®^). Ultra-running led to oxidative stress, metabolic shifts, and inflammation-driven RBC damage, including increased acylcarnitines, kynurenine accumulation, oxidative lipid and protein modifications, reduced RBC deformability, enhanced microparticle release, and increased senescence markers such as externalized phosphatidylserine (PS). Post-race interleukin-6 strongly correlated with kynurenine elevation, mirroring inflammatory responses in severe infections. These findings challenge the assumption that RBC damage in endurance exercise is primarily mechanical, revealing systemic inflammation and metabolic remodeling as key drivers. This study underscores RBCs as both mediators and casualties of extreme exercise stress, with implications for optimizing athlete recovery, endurance training, and understanding inflammation-linked RBC dysfunction in clinical settings.

**Teaser:** Marathon running imparts molecular damage to red blood cells, the effects of which are exacerbated by increased distances of ultramarathons.

## Introduction

Endurance exercise places significant demands on the oxygen transport system, and red blood cells (RBCs) are at the forefront of meeting these challenges. In addition to delivering oxygen and removing carbon dioxide, RBCs contribute to pH buffering and release signaling molecules— such as ATP and nitric oxide—that facilitate vasodilation and enhance muscle perfusion (*1, 2*). While moderate exercise is associated with improved cardiovascular function and metabolic health, prolonged endurance events, like marathons and ultramarathons, impose additional mechanical, metabolic, and oxidative stresses. These stressors can result in acute cellular damage particularly in RBCs, where the capacity for repair is limited owing to the lack of nuclei and organelles that results in the incapacity to replace damaged protein and lipid components through *de novo* synthesis (*3*).

Historically, investigations into exercise physiology have largely focused on skeletal muscle (*4*), serum (*5*), and plasma (*6*) biomarkers to gauge the systemic impact of prolonged running. However, emerging evidence suggests that the RBC compartment provides critical insights into the cellular adaptations and injury responses elicited by extreme exercise (*7, 8*). Yet, classic concepts like foot-strike hemolysis – also known as march hemoglobinuria, first described by Fleischer in 1881 (*9*) – have been recently challenged in comprehensive reviews (*10*), which indicates an incomplete understanding of the role of RBC biology in exercise tolerance and performance. Recent advances in multiomics technologies have opened new avenues for comprehensively mapping the molecular responses to strenuous exercise. Simultaneous interrogation of the metabolome, lipidome, proteome, and metallome, enables holistic understanding of how tissues and circulating cells respond to exercise-induced stress. Studies have highlighted that transient cytokine release (e.g., interleukin-6) during endurance exercise triggers metabolic remodeling and tissue repair (*11, 12*), in addition to oxidative modifications in RBC, shifts in membrane lipid composition, and ultimately compromised RBC deformability (*7*).

Despite these advances, the specific molecular responses of RBCs to prolonged endurance exercise remain insufficiently characterized. This knowledge gap is particularly relevant in the context of long-distance trail running, where additional challenges such as variable terrain, altitude, and prolonged mechanical stress further compound the injury and repair processes in RBCs. Indeed, long distance running results in appreciable damage to RBC including decreased deformability and increased markers of senescence (*13, 14*) in the face of increased inflammation (*15*). Previous omics-based studies have predominantly focused on plasma responses to acute exercise or short-duration laboratory tests (*16*), ultramarathon treadmill simulations (*17*), or *in situ* (*18, 19*), which cannot recapitulate the unique stressors encountered during ultra-endurance events in the outdoors [reviewed in (*20*)]. Moreover, while plasma-based analyses provide valuable systemic information, they do not capture the direct impact on RBCs—cells that lack nuclei and organelles and are thus uniquely dependent on metabolic pathways to cope with oxidative damage.

In light of these considerations, our study was designed to integrate multi-omics approaches to delineate the impact of long-distance trail running on both plasma and RBC molecular signatures. By comparing pre- and post-race samples from athletes competing in the World Class Martigny– Combes à Chamonix race (MCC, 40 km, elevation gain: 2,300 m) and Ultra-Trail du Mont Blanc race (UTMB^©^, 171 km, elevation gain: 10,000 m), we aimed to elucidate the distinct metabolic, inflammatory, and oxidative modifications that occur with increasing exercise duration. We hypothesized that while both marathon and ultramarathon events trigger systemic inflammatory responses and metabolic remodeling, the extent of cellular damage—particularly within the RBC compartment—would be more pronounced in ultramarathon running.

## Results

### Molecular responses to ultra-running in plasma and RBCs

To characterize molecular changes in both plasma and RBCs as a function of running distance, samples were isolated from 23 runners, divided as follows: (i) 11 runners (5 female, 6 male, 35.7 ± 8.6 years) before (Pre) and immediately after (Post) the MCC; or (ii) 12 runners (4 female, 8 male, 38 ± 6.4 years) Pre and Post UTMB. In addition to assessing hematological and rheological parameters (*14*), mass spectrometry-based metabolomics, lipidomics, trace element analysis (i.e. metallomics), and proteomics profiles were determined (**Figure 1A**). After curation, these analyses provided quantitative data in plasma or RBCs on 440 or 1105 proteins, 659 or 647 lipids, 197 or 271 metabolites, and 8 or 6 metals, respectively (**Figure 1B, Supplementary Tables 1 and 2**, respectively). Using the entire dataset, Partial Least Squares-Discriminant Analysis (PLS-DA) separated Pre from Post plasma samples along Component 1, which explained 11.1% or 17.5% of the data variance in the MCC or UTMB, respectively (**Figure 1C-D**). While MCC plasma was largely characterized by increases in acyl-carnitines (AC) and fatty acids (FA), UTMB plasma showed decreased levels of lysophospholipids (LP) and increases in the purine catabolite, hypoxanthine, muscle creatine kinase (KCRM), and acute phase response proteins CRP, SAA1, SAA2 (**Figure 1C-D**). Similarly, RBCs after the MCC had increased AC (including hydroxylated), FA, and decreased coenzyme A precursor pantothenate, while RBCs after the UTMB had elevated levels of (hydroxy)AC, the oxylipin 13(S)-HODE, and tryptophan catabolite, kynurenine (**Figure 1E-F**). Mean running speed was significantly faster (Welch’s t-test p = 0.009) in the MCC than the UTMB at 6.07 ± 1.61 km/hr or 4.43 ± 0.92 km/hr, respectively (**Figure 1G**). However, a linear mixed model of run speed (adjusted for age, sex, and BMI) using fold-change values (Post/Pre) showed significant associations with 51 plasma molecules that largely included lipids such as phosphatidylcholines (PC), acylcarnitines (AC), cholesteryl esters (ChE), ceramides (Cer), sphingomyelins (SM), diacylglycerides (DG), and triacylglycerides (TG), without or with ether (O-) linkages (**Figure 1H**). This model also highlighted tryptophan metabolism (kynurenine, kynurenate, indolepyruvate), and transferrin receptor (TFR1), which mediates cellular iron uptake. Meanwhile, only the RBC level of adenosine diphosphate (ADP) was significantly associated with run speed (**Figure 1I**).

**Figure 1.**
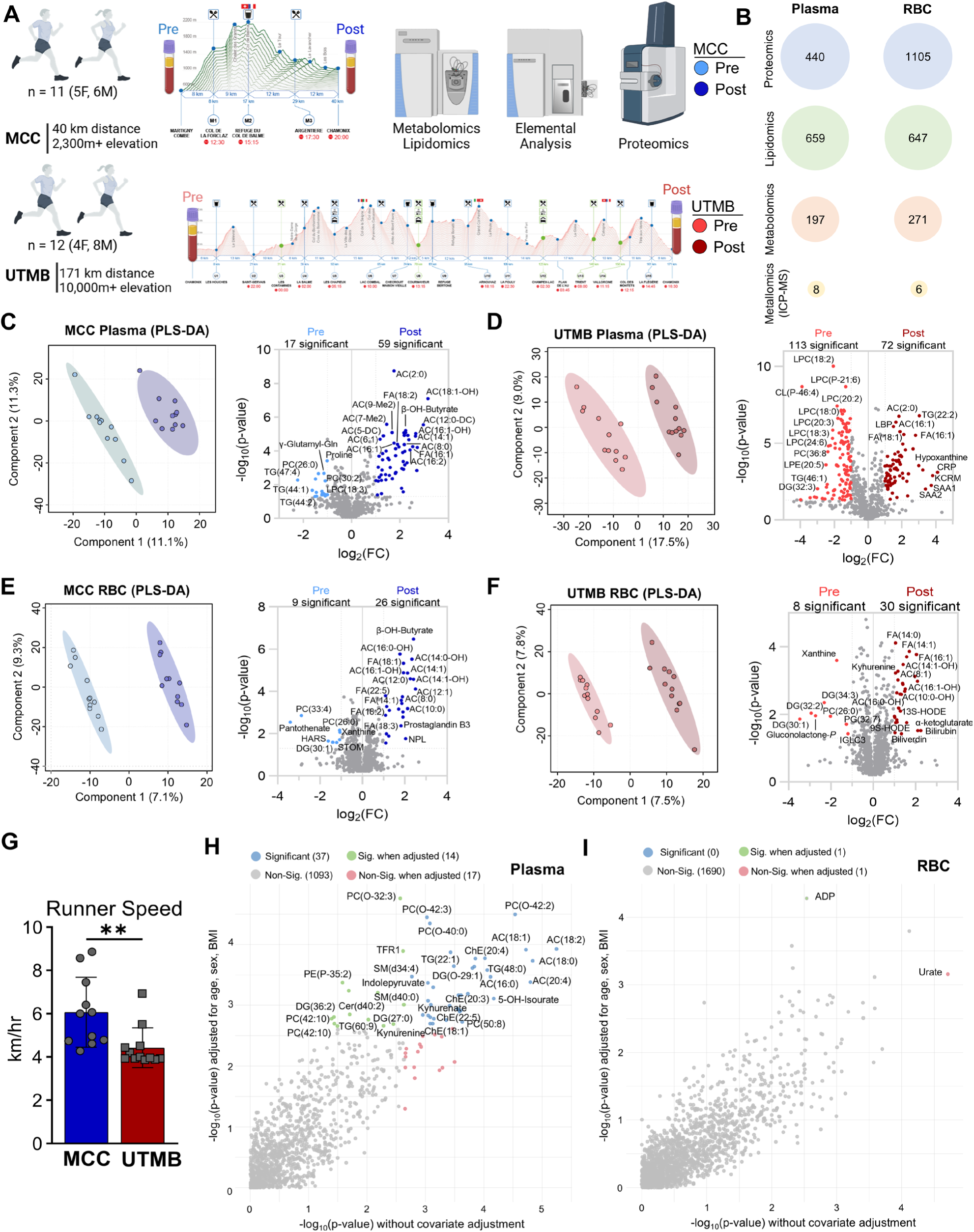
Multiomics of a 40 km or 171 km Run. (A) Plasma and RBC were taken before (Pre) and after (Post) a 40-km (Martigny–Combes à Chamonix race, MCC) or 141-km (Ultra-Trail du Mont Blanc, UTMB^©^) footrace. (B) Proteomics, lipidomics, metabolomics coupled plasmon (ICP)-MS were collected. Processed data contained measurements for the number of analytes shown in the respective circle for each fraction. (C) Partial Least Squares Discriminant Analysis (PLS-DA) of the plasma fraction during the MCC and (D) UTMB along with associated volcano plots highlighting all analytes with fold change (FC) > 1.5 and p-value < 0.05 from a paired T-test. (E) PLS-DA of the RBC fraction during the MCC and (F) UTMB along with associated volcano plots highlighting all analytes with fold change (FC) > 1.5 and p-value < 0.05 from a paired T-test. (G) Average speed (in kilometers per hour) for all runners in each race. (H) Linear mixed model adjusted for age, sex, BMI of runner speed (km/h) in the plasma, or (I) RBC fraction.

### Inflammatory markers increase in the plasma after ultra-running

Considering that molecular signatures in the plasma were more strongly associated with running time, we next identified signatures with respect to individual molecular class (i.e. proteins, metabolites, lipids). Reflecting the strong lipidomic signature associated with both speed and distance, MCC and UTMB runners’ plasma had significantly decreased diacylglycerides (DG), while UTMB alone also had significant decreases in lysophosphatidylcholines (LPC), lysophosphatidylethanolamines (LPE), and cardiolipins (CL), alongside increased acylcarnitines (**Figure 2A**). Active lipid metabolic pathways during both MCC and UTMB were mapped using Bioinformatics Methodology For Pathway Analysis (BioPAN) (*21*) and revealed distinct lipid remodeling patterns between the two races (**Figure 2B, Supplementary Table 3**). In UTMB, the lysophosphatidylethanolamine (LPE) → phosphatidylethanolamine (PE) → phosphatidylserine (PS) axis was significantly enriched, suggesting an increased demand for PE and PS. In contrast, MCC exhibited a stronger conversion of PE to phosphatidylcholine (PC) likely driven by phosphatidylethanolamine N-methyltransferase (PEMT) activity, indicating a preference for PC biosynthesis over PS production. Additionally, UTMB showed a pronounced increase in ether lipid remodeling (O-LPC → O-PC), while MCC displayed a broader remodeling of glycerophospholipids, particularly in PC and LPS pathways. These differences highlight distinct metabolic adaptations between the two conditions, reflecting variations in membrane composition,

**Figure 2.**
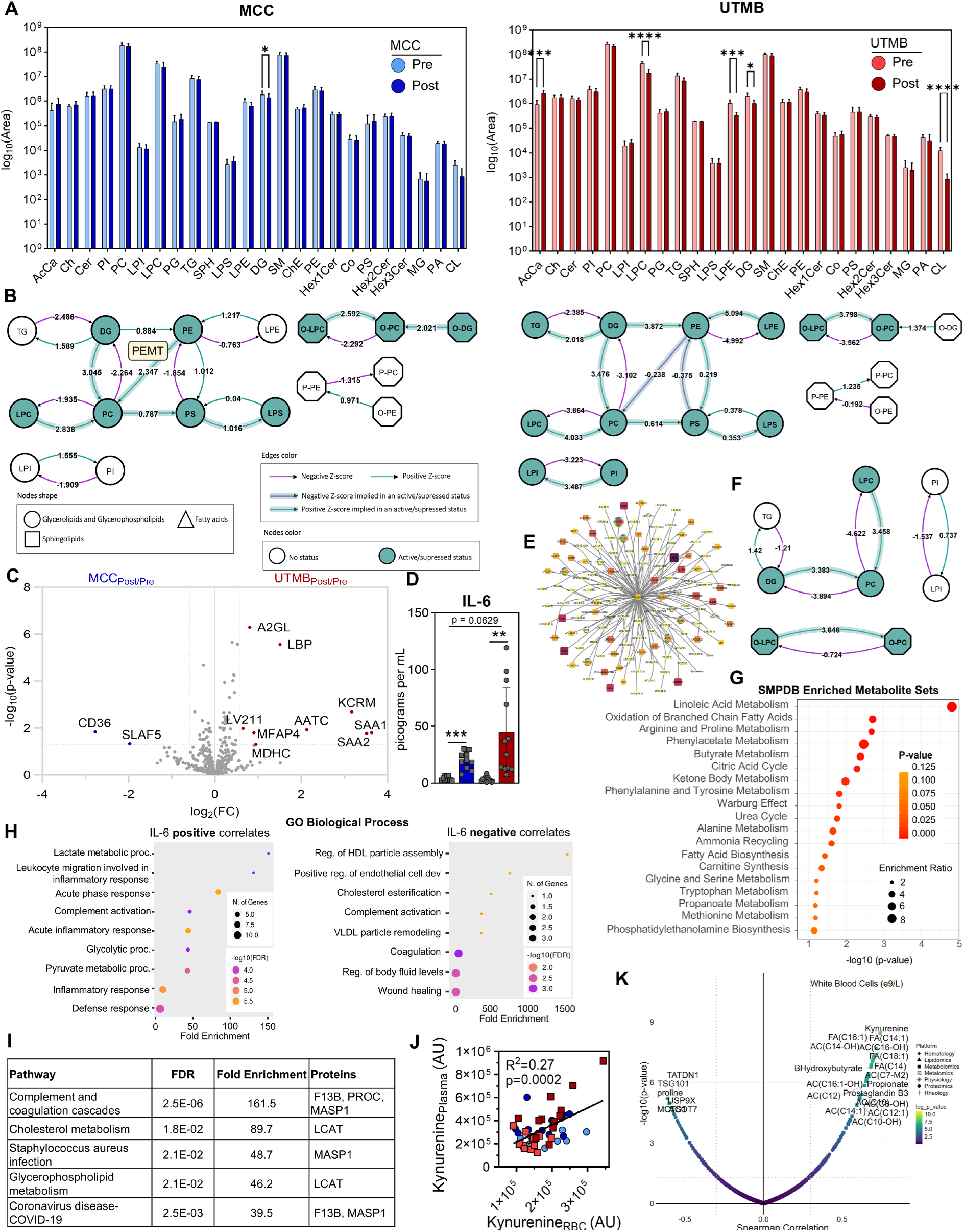
Lipidomics of the MCC and UTMB. (A) Total lipid class quantities shown as log_10_(Peak area sum) before and after the MCC and UTMB. (B) BioPAN models of lipid class flux during the MCC (left) and UTMB (right). Lipid classes are indicated by node shape where filled nodes indicate significantly modified class. Fluxes normalized by Z-score with positive fluxes shown in teal and negative fluxes shown in purple. (C) A volcano plot showing the intra-subject proteomic fold changes with respect to each race. (D) IL-6 measurements before and after the MCC (blue) and UTMB (red). (E) A Debiased Sparse Supplementary Figure 2. (F) BioPAN model of lipid pathway flux for lipids significantly associated with IL-6 levels. (G) Metabolite set enrichment analysis for metabolites lipids significantly associated with IL-6 levels. (H) Gene Ontology analyses for proteins positively (left) of negatively (right) significantly associated with IL-6 levels. (I) KEGG pathway enrichment based on proteomics data. (J) Kynurenine correlations between plasma (y-axis) and RBC (x-axis) compartment. (K) RBC molecular correlates with plasma IL-6 levels.

Changes in the plasma proteome contained fewer significant differences, although the number of significantly increased proteins after the race was higher in the UTMB cohort (**Supplementary Figure 1**). When comparing fold changes at completion of each race to baseline values, and consistent with the race-specific lipidomics signatures noted above, post-race elevation of the fatty acid transporter and immunomodulator CD36 (*22*) was observed particularly in MCC runners (**Figure 2C**). On the other hand, UTMB runners presented with larger increases in creatine kinase muscle-type (KCRM), which is indicative of ongoing skeletal muscle damage (*23*) (**Figure 2C**). In addition, compared to MCC, UTMB runners had elevations in numerous proteins involved in the acute phase response including Serum Amyloid A1 and A2 (SAA1 and SAA2) and LPS-binding protein (LBP), which plays a critical role in the innate immune response to bacterial infections. In support of these proteomics data, the circulating levels of multiple cytokines (*15*) including IL-6 were significantly higher in runners after both races, with a trend towards higher elevations after the UTMB (**Figure 2D**). As such, a Debiased Sparse Partial Correlation (DSPC) network (*24*) of IL-6 revealed several associations with total white blood cell counts, inflammatory proteins (CRP, SAA1, SAA2, SAA4, LBP, A2GL, A1AG1), cytosolic proteins (TKT, MDHC, LDHB), carboxylic acids (citrate, succinate, fumarate, malate), purine catabolites (hypoxanthine, urate), acylcarnitines, and fatty acids (**Figure 2E, Supplementary Figure 2**). Lipid pathways that correlated with IL-6 levels centered largely upon DG and LPC conversion into PC (**Figure 2F**). Plasma metabolites that correlated with IL-6 levels were significantly involved in linoleic acid metabolism, though there was substantial representation from mitochondrial pathways including the Citric Acid Cycle, Urea Cycle, branched chain amino acid oxidation, and metabolism of many amino acids (**Figure 2G**). Gene ontology (GO) analysis of proteins that were positively correlated with IL-6 confirmed acute inflammatory response, while downregulated proteins were involved in cholesterol trafficking (lipoprotein particle assembly), complement activation, coagulation, regulation of body fluid levels, and wound healing (**Figure 2H**). Interestingly, negative correlated proteins F13B and MASP1 have also been associated with COVID-19 pathology (**Figure 2I**). Kynurenine was an additional molecular correlate with IL-6 (**Figure 2E, Supplementary Figure 2**), also consistent with this reportedly strong positive association in COVID-19 disease severity.(*25–27*) While kynurenine levels significantly correlated between the plasma and RBCs fractions in this study (**Figure 2J**), RBC kynurenine levels in particular were the strongest molecular correlate with IL-6 across the entire dataset (**Figure 2K, Supplementary Figures 3-4**), consistent with the hypothesis that uptake via SLC7A5 (LAT1) favors a role for RBCs as a circulating reservoir of metabolic markers of systemic inflammation.(*28*)

### RBCs accumulate damage markers specifically during ultra-running

In light of the strong link between RBC molecular signatures and inflammation, and the recently reported association between elevated RBC kynurenine levels and increased susceptibility to hemolysis (*28*), we next turned to in depth analysis of the RBC fraction. Multiple changes to RBC characteristics were distinctly observed after running a marathon or ultra-marathon distance (**Figure 3A**). Namely, mean cell volume (MCV) decreased after both races in line with increases in immature reticulocytes. While elevations in PS-exposing RBCs (a marker of eryptosis (*29–31*)) were observed after the MCC, PS-exposing RBCs were unchanged after the UTMB. However, UTMB runners showed an increase in circulating RBC-derived microparticles in addition to an appreciable reduction in hematocrit. To determine molecular changes to RBCs as they pertain to running distance, PLS-DA using fold change data (Delta = Post/Pre) was able to distinguish RBCs from each race (**Figure 3B**). The molecules that most strongly influenced this clustering pattern included urate, creatinine, medium chain acyl-carnitines, and LPCs – higher after MCC – or long chain acylcarnitines, carboxylic acids (fumarate, methylmalonate), carnitine, and copper – higher after UTMB (**Figure 3C**).

**Figure 3.**
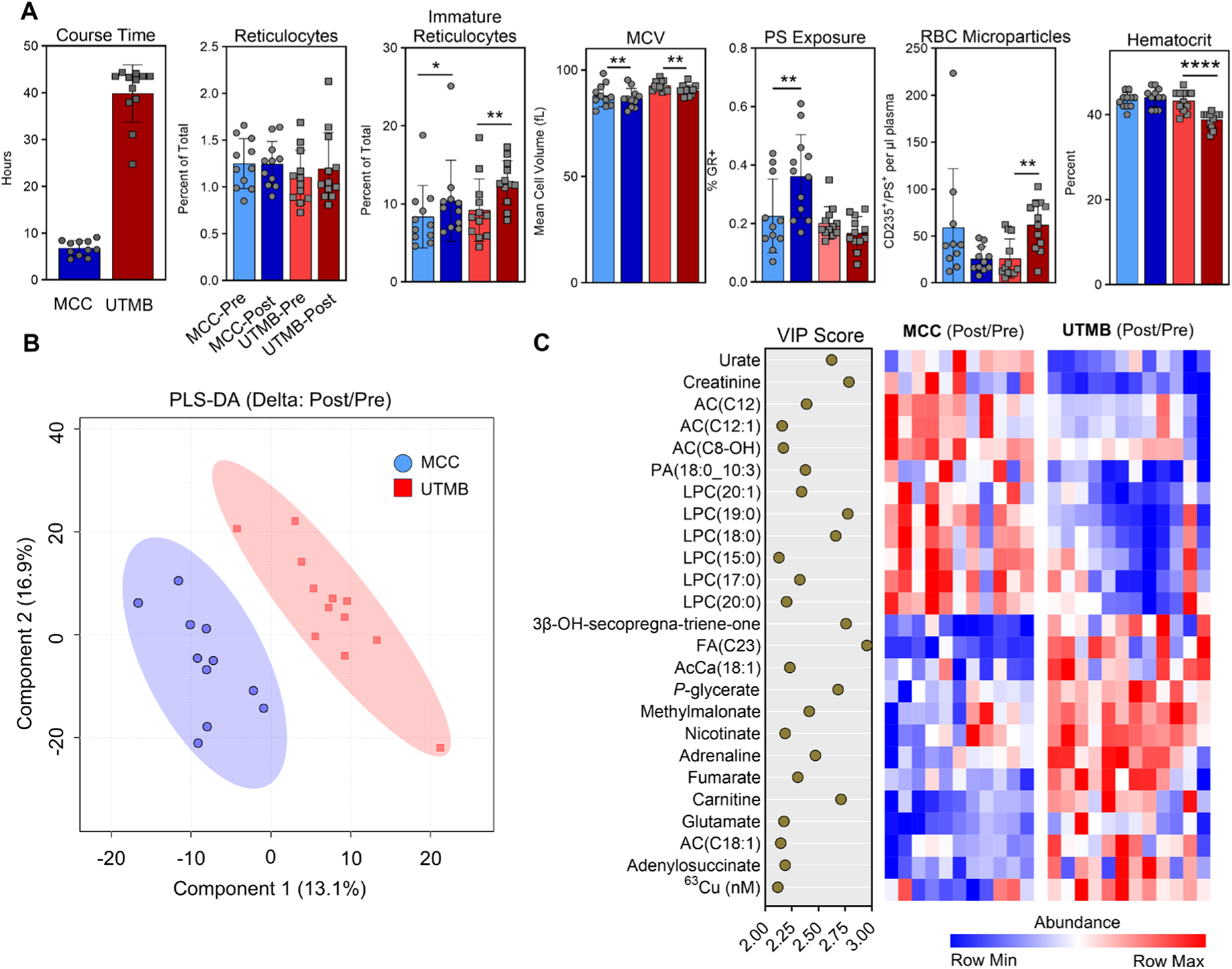
RBC Multiomics of Running. (A) Hematologic parameters of each race is shown. (B) PLS-DA of intra-subject normalized molecular values for each race. (C) The top 25 molecules based on Variable Importance in Projection (VIP) for the PLS-DA are shown (left) along with relative values of intra-subject fold changes for each race in the heat map (right).

### RBCs activate Lands Cycle for membrane lipid repair during marathon and ultra-marathon running

Changes in the abundance of multiple lipid classes as a whole occurred during both races (**Figure 4A**). Notably, acylcarnitines were significantly elevated after the UTMB in conjunction with decreased LPC. In RBCs, these lipid classes are central to the Lands Cycle, a process by which RBCs replace oxidatively damaged membrane lipid acyl chains using a pool of acyl-CoA molecules that are interconverted with acylcarnitines based on demand for lipid repair (**Figure 4B**). Interrogation of specific free fatty acids (**Figure 4C**) and acylcarnitines (**Figure 4D**) highlighted increased levels after each race. In line with an activated Lands cycle, the ratio of lysophospholipids-to-phospholipids (specifically LPC/PC and LPE/PE) were lower after each race (**Figure 4E**). The extent of both fatty acid and medium chain acylcarnitine accumulation was higher in RBCs after the MCC, while post-UTMB RBCs maintained comparable levels of long chain acylcarnitines including the most abundant AC(18:1). RBCs rely on pantothenate to synthesize CoA, which significantly decreased after both races. On the other hand, carnitine was significantly elevated after the UTMB (**Figure 4F**), suggesting decreased utilization for synthesis of acylcarnitines in these RBCs.

**Figure 4.**
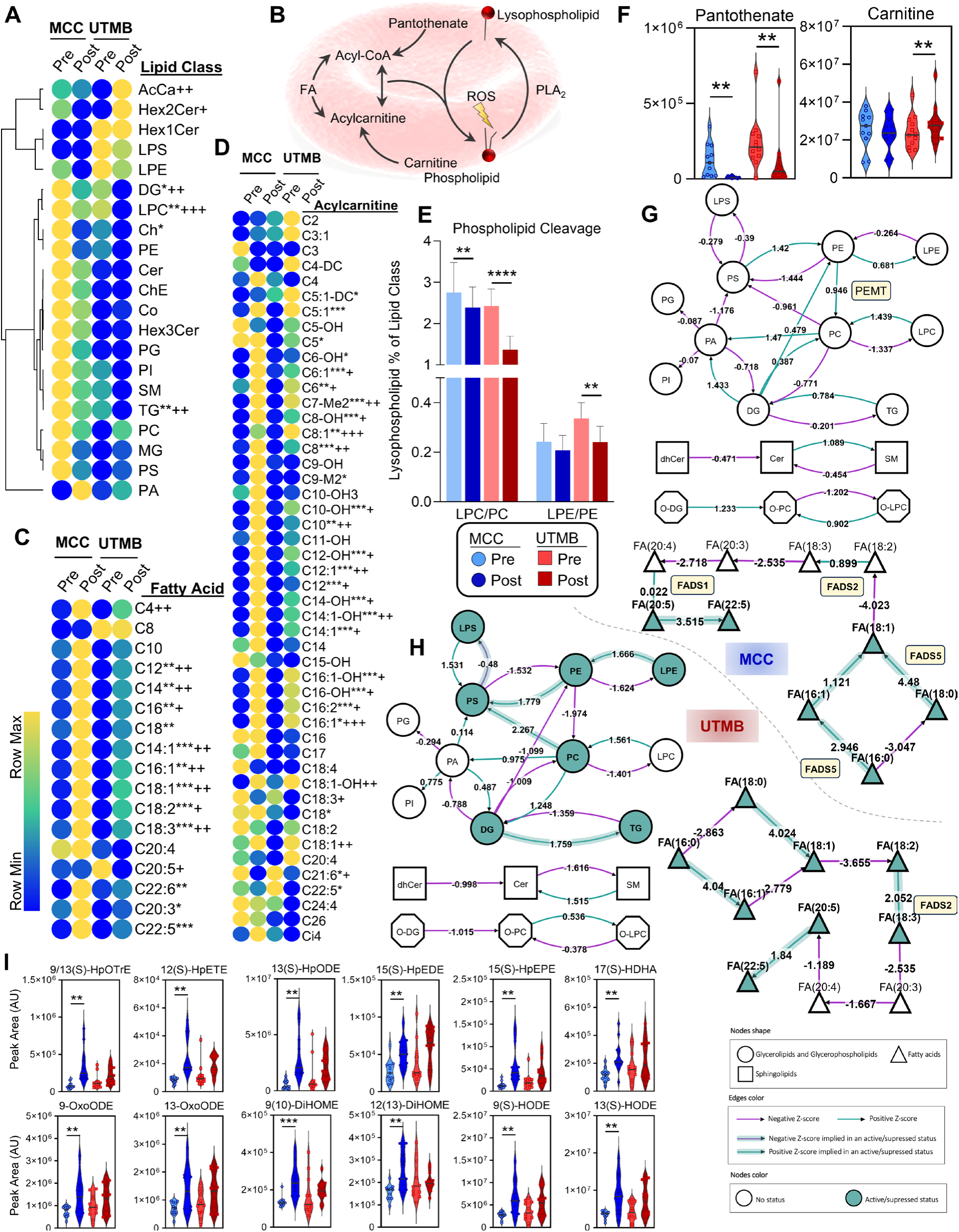
RBC Lipidomics. (A) The average peak areas of lipids identified from untargeted analysis were summed according to lipid class and plotted as a heat map. p-values from Two-tailed paired T-test for MCC pre/post comparisons are indicated as *, p<0.05; **, p<0.01, ***, p<0.001; ****, p<0.0001 and p-values for UTMB pre/post comparisons are indicated as +, p<0.05; ++, p<0.01, +++, p<0.001; ++++, p<0.0001. (B) A cartoon of the Lands cycle for red blood cell membrane repair is shown. (C) Average peak areas from semi-targeted analysis for fatty acids and (D) acylcarnitines are shown as heat maps. (E) levels of pantothenate and carnitine are plotted as violin plots. (G) BioPAN or RBC during the MCC and (H) UTMB are shown. (I) Violin plots of significant oxylipins before and after the MCC (blue) and UTMB (red).

Lipid network analysis using BioPAN revealed distinct lipid remodeling patterns in RBCs between MCC and UTMB (**Figure 4G and 4H, Supplementary Table 4**). In MCC, RBCs exhibited a stronger PE → PC remodeling pathway mediated by PEMT. Additionally, MCC showed significant ether lipid remodeling (O-LPC → O-PC), driven by Lysophosphatidylcholine Acyltransferase 3 (LPCAT3), which is linked to Lands cycle activation, ferroptosis and extravascular hemolysis (*32*). This pattern suggests increased oxidative stress, as ether lipids play a crucial role in protecting RBCs membranes from lipid peroxidation (*33*). In contrast, UTMB RBCs exhibited a stronger shift toward phosphatidylserine (PS), with significant activity in PC → PS (Z = 2.267) and PE → PS (Z = 1.779) conversions that are mediated by PTDSS1 and PTDSS2 (Phosphatidylserine Synthases 1 & 2). These reactions, along with the LPE → PE → PS → LPS pathway, indicate a preference for redistributing phospholipids within the membrane. However, there was no corresponding increase in PS externalization (**Figure 3A**), suggesting either that the population of RBCs in circulation after the UTMB maintained membrane asymmetry as a function of increased ATP-dependent flippase activity, or that RBCs with exposed PS were removed from circulation during the course of the UTMB.

Distinct differences between MCC and UTMB in fatty acid homeostasis was observed as well (**Figure 4G** and **4H**). RBCs from both races displayed increased fatty acid desaturation activity – conversion of FA(16:0) → FA(16:1) and FA(18:0) → FA(18:1) via FADS5. However, MCC RBCs demonstrated increased FA(18:2) → FA(18:3) conversion (Z = 0.899), while this conversion was notably larger (Z = 2.052) in UTMB RBCs. As this step is carried out by Fatty Acid Desaturase 2 (FADS2), which is present in RBCs and active in maintaining NAD/NADH ratios under oxidant stress (*34*), differences between RBCs during MCC and UTMB suggest altered networks to maintain redox homeostasis in the face of ongoing endurance running.

Additionally, eicosanoid and oxidized lipid species showed differential regulation between MCC and UTMB (**Figure 4I**). While HETEs (hydroxyeicosatetraenoic acids) remained unchanged (**Supplementary Table 2**), significant (p < 0.05) increases were observed in MCC of several hydroxy derivatives (including 9(S)- and 13(S)-HODE), hydroperoxy derivatives (including 9(S)-HpOTrE, 12(S)-HpETE, 15(S)-HpEDE, 9(S)- and 13(S)-HpODE, and 15(S)-HpEPE), which are formed by lipoxygenase-mediated or non-enzymatic auto-oxidation of polyunsaturated fatty acids. Moreover, increases in downstream metabolites, such as oxo forms (including 9(S)- and 13(S)-oxoODE) and dihydroxy forms (including 9(10)- and 12(13)-diHOME), indicate significantly increased lipid peroxidation and oxidative stress in MCC RBC, while these molecules approach significance (p < 0.08) in UTMB RBC.

### Prolonged running activates purine salvage and alternative carboxylate metabolism

Because they are devoid of mitochondria, RBCs depend solely on glycolysis for ATP production. RBCs after the MCC were not different aside from significantly accumulated lactate (**Supplementary Figure 5**). However, the UTMB elicited significant accumulations in multiple glycolytic intermediates aside from pyruvate and lactate that trended higher, but did not reach significance. Increased phosphoenolpyruvate/pyruvate ratios in UTMB, but not MCC, is consistent with inhibition of redox sensitive pyruvate kinase (PK). However, it remains to be assessed whether pharmacological PK activation could be beneficial in the context of an ultra-marathon, in that PK activation would come at the cost of 2,3-BPG depletion (*35*) thus counteracting the beneficial effects this molecule has on hemoglobin allostery by favoring O_2_ off-loading. Similarly, RBCs after the UTMB had lower oxidative phase Pentose Phosphate Pathway (PPP) intermediates and higher end-products (ribose phosphate and pentose phosphate isomers), including downstream non-oxidative phase sedoheptulose phosphate, which is suggestive of an active flux through this pathway.

The PPP is used to generate NADPH for oxidative stress management, which impacts purine levels that are maintained through the purine salvage pathway (*36*). RBCs after the MCC had higher inosine and urate, indicative of purine catabolism (**Figure 5A**). In contrast, RBCs after the UTMB had higher IMP and hypoxanthine, which is derived from oxidant stress activation of AMPD3 to trigger purine breakdown and deamination (*37*). Subsequent decreases in xanthine and allantoate are further indicative of purine salvage rather than catabolism. This pathway relies upon aspartate, which is used by adenylosuccinate synthase (ADSS) to convert IMP back into AMP. The metabolic signature of carboxylic acids in UTMB RBCs demonstrates increased utilization of aspartate in this process (**Figure 5B**). In addition, elevated levels of malate suggest activation of the GOT1/MDH axis to recycle NAD^+^, in similar fashion to previous studies of RBCs during acclimatization to high altitude or hypoxic storage in the blood bank (*38*). Similarly, pyruvate/lactate ratios were higher both at baseline and after the race in UTMB compared to MCC (**Figure 5C**), suggestive of the activation of methemoglobin (MetHb) reductase. Indeed, as LDH and MetHb reductase compete for NADH as a substrate, elevation in pyruvate/lactate ratios reflect oxidant metabolic defects in RBCs, such as in the case of G6PD deficiency (*39, 40*).

**Figure 5.**
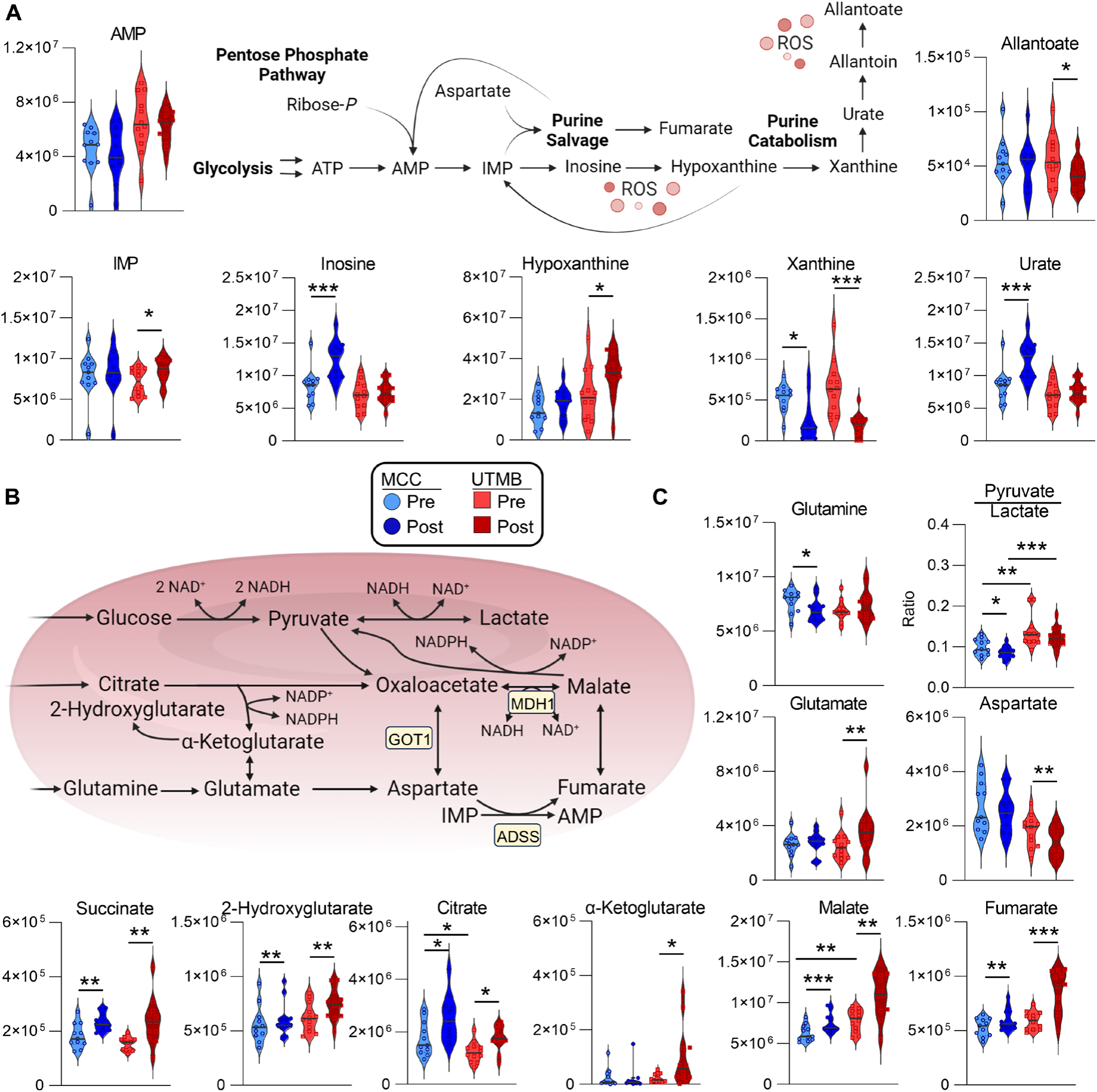
Purine Salvage and Carboxylate Metabolism in RBC. Metabolites in the (A) purine salvage or (B) carboxylate metabolism pathways are shown. Individual samples from MCC (O) at Pre (light blue) and Post (dark blue), or UTMB (Δ) at Pre (light red) or Post (dark red) are shown within violin plots. p-values from Two-tailed homoscedastic T-tests are indicated as *, p<0.05; **, p<0.01, ***, p<0.001; ****, p<0.0001. The pathway map (center) was created with Biorender.com.

### The RBC proteome accumulates oxidative damage during marathon and ultra-marathon running

After establishing the impact of both MCC and UTMB – especially the latter – in triggering oxidant stress to RBCs, we then sought to determine the impact of running distance on protein homeostasis, given the incapacity of mature RBCs to replace oxidatively damaged proteins via *de novo* protein synthesis and the concurrent reliance of aging RBCs on proteasomal machinery to remove damaged proteins (*41*). To this end, RBC proteomics data were mined for redox-related post-translation modifications (PTM) including methionine oxidation, asparagine deamidation, and glutamate/aspartate methylation, all of which are marks of protein damage and repair in RBCs (*42*). Unsupervised principal component analysis (PCA) using peptide level data (including PTMs) was capable of distinguishing Pre from Post samples in both races across component 1, which described 44.3% of the data variance (**Figure 6A, Supplementary Table 5**). Of the multiple PTMs detected across the proteome, only methionine oxidation was significantly elevated after both the MCC and UTMB (**Figure 6B**). Correlating total PTM levels with hematological and blood rheological parameters including plasma viscosity, RBC deformability, microparticle generation, and cell volume revealed that methionine oxidation had the largest number of positive correlates and carbamidomethylated cysteine – a marker of reduced thiol content – had the largest number of negative correlations (**Figure 6C**). Total carbamidomethyl cysteine content was negatively (R < 0.75) associated with runner body fat percentage and IL-6 levels (**Figure 6C**). In parallel, specific parameters including blood viscosity and RBC deformability (EI – elongation index) as well as runner age, while the top negative correlate was the reduced (GSH) to oxidized (GSSG) glutathione ratio (**Figure 6C**). Furthermore, the top 100 peptides by ANOVA exclusively contained methionine oxidation (**Figure 6D**). RBCs after the UTMB, however, contained a higher degree of methionine oxidation when looking at modified peptide as a function of total peptide signal (**Figure 6E**), thereby indicating an increased degree of proteome oxidation in these samples. KEGG pathway analysis of proteins with statistically significant (ANOVA FDR < 0.05) increases in methionine oxidation revealed a substantial impact to components within the proteasome, Pentose Phosphate Pathway, and redox proteostasis including cysteine, methionine, and glutathione metabolism (**Figure 6F, Supplemental Figure 6**). Multiple neurodegenerative pathways were also enriched including Spinocerebellar ataxia, Parkinsons disease, Huntington disease, Alzheimer disease, and Amyotrophic lateral sclerosis. These pathways largely share proteasome, redox, and cytoskeletal proteins (**Supplementary Figure 7B**). Collectively, these analyses indicate an enrichment in four main categories of proteins, rather than random oxidation events:

1. Antioxidant enzymes: PRDX6, PCMT1 – which participates in isoaspartyl damage repair (*42*) – a modification that was elevated in MCC RBCs after the race – (**Figure 6B)**; GCLM/GSTO - linked to GSH synthesis and utilization; HPRT1 - linked to purine salvage; LTA4H - linked to eicosanoid metabolism;
2. Metabolic enzymes: BPGM, PGD, BLVRB, ACLY – explaining in part the observed changes in 2,3-BPG levels, PPP and CoA homeostasis and citrate levels – **Supplementary Figure 5**);
3. Ubiquitination systems and proteasome components: UBA1, UBE2V2, PSMC3; in the context of the ubiquitin system, the active site cystine within E1 and E2 enzymes forms a thioester bond with ubiquitin, facilitating its activation and transfer for ubiquitination (*43*); however, specific searches for post-translational markers of ubiquitination (K-GG modification) showed no significant changes between baseline and completion in either of the two races – (**Figure 6B)**;
4. Structural proteins: SPTA1, SPTB, ANK1, SLC4A1, DMTN, STOM, ADD – consistent with elevated microparticle release (**Figure 3A**) and suggestive of impaired RBC deformability.

**Figure 6.**
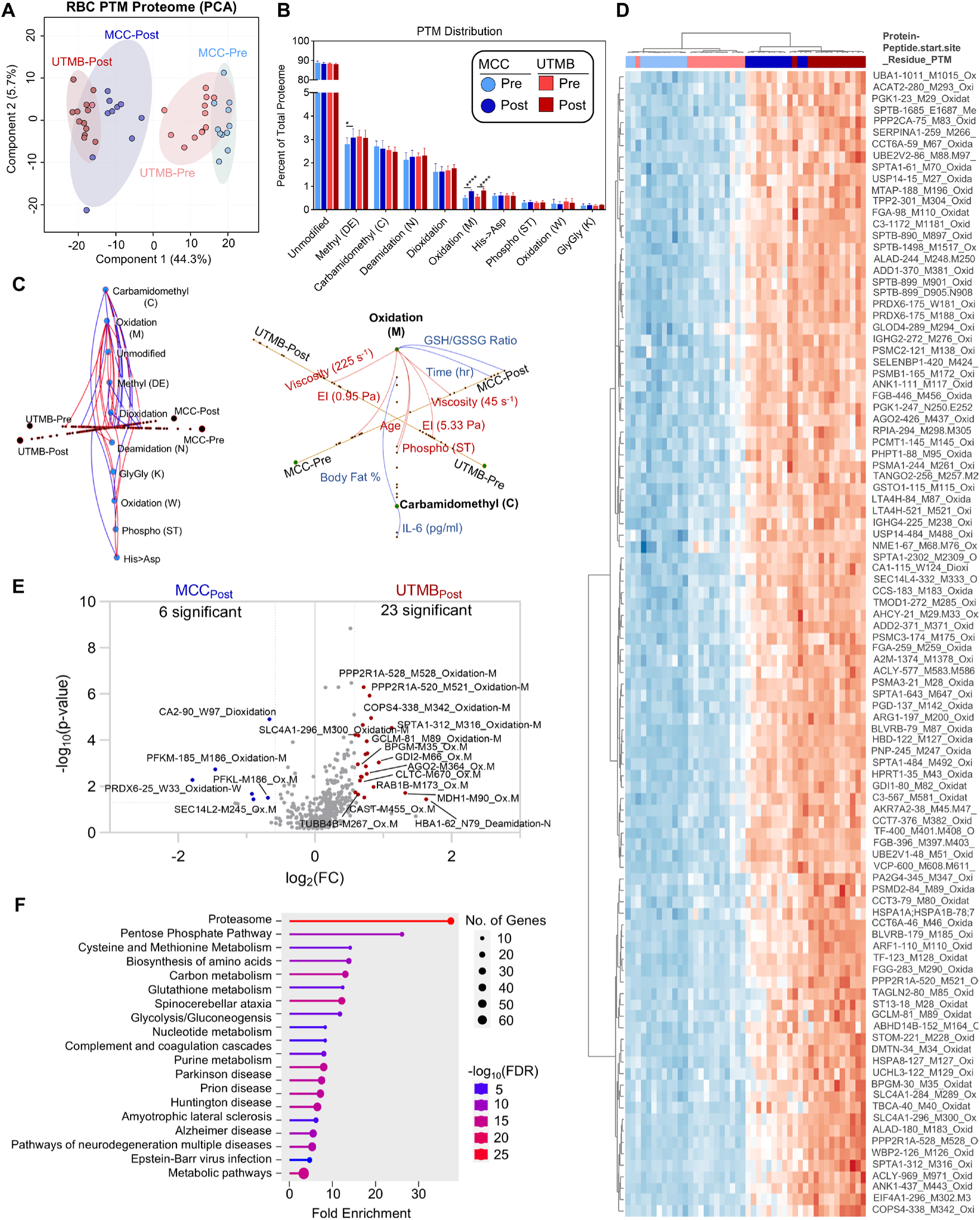
RBC Proteomics demonstrates increased protein oxidation. (A) Principal Component Analysis (PCA) of RBC peptide-level data. (B) Distribution of post translational modification (PTM) across the proteome. (C) Hive plot of significant Pearson correlations (R) between physiological, hematological, and rheological parameters are shown with R > 0.7 shown in red and R < −0.7 shown in blue. On the right, a labeled plot showing Pearson correlations (R > 0.7, R < −0.7) with methionine oxidation and cysteine carbamidomethylation. (D) Heat map showing relative peptide abundance across MCC and UTMB. (E) Volcano plot comparing peptide levels in the post MCC and UTMB samples. (F) KEGG Pathway enrichment of proteins with statistically significant (ANOVA Fisher’s LSD FDR < 0.05) enrichment of methionine oxidation.

### RBC deformability correlates with protein homeostasis and copper

Given the significant accumulations of lipid and protein oxidation, inflammatory response (specifically IL-6 and kynurenine), and decreased hematocrit after the UTMB, we next focused on blood rheological parameters. Hematological and blood rheological parameters were assessed before and after each race (*14*) including blood viscosity and RBC deformability (**Supplementary Figure 8**). While blood viscosity showed variable changes (increasing at 11.5 cP/s after MCC and decreasing at 225 cP/s after UTMB), RBC deformability was markedly decreased at higher shear stresses specifically after UTMB. Integration of the area under the shear stress vs EI curve for each participant enabled calculation of a single RBC deformability value, which significantly decreased after the UTMB (**Figure 7A**). This approach also allowed for correlation of all multi-omics measurements with composite RBC deformability and revealed significant positive correlations with metabolites involved in nitrogen metabolism (lysine, proline, xanthine, ornithine), while glycolytic intermediates (2/3-phosphoglycerate, 2,3-BPG) and hydroxyacyl-carnitines including AC(14:0-OH), AC(14:1-OH) and AC(16:1-OH) were significant negative correlates (**Figure 7B**). GO enrichment of significantly correlated proteins highlighted proteasome assembly as the most enriched term (**Figure 7C, top**). Despite significantly correlating with RBC deformability, PSMA2, either MCC or UTMB (**Figure 7D**). However, when factoring in PTM status on proteasome subunits, increased methionine oxidation (node size is proportional to average fold change for both races) inversely correlated with RBC deformability (**Figure 7E**). Therefore, these results implicate accumulating proteasome oxidation that could impair its activity to remove damaged proteins with decreasing RBC deformability. Additional correlations with RBC deformability were observed with metals (**Figure 7F**). Potassium, magnesium, and iron had weak positive correlations with EI at lower shear stress, though these correlations were lost (or inverted in the case of iron) at shear stresses greater than 10 Pa. Most notably, copper levels were inversely correlated with RBC deformability at all shear stresses. The concentration of copper, calcium, potassium, iron, magnesium, and sodium were elevated in RBCs specifically after the UTMB (**Figure 6G**).

**Figure 7.**
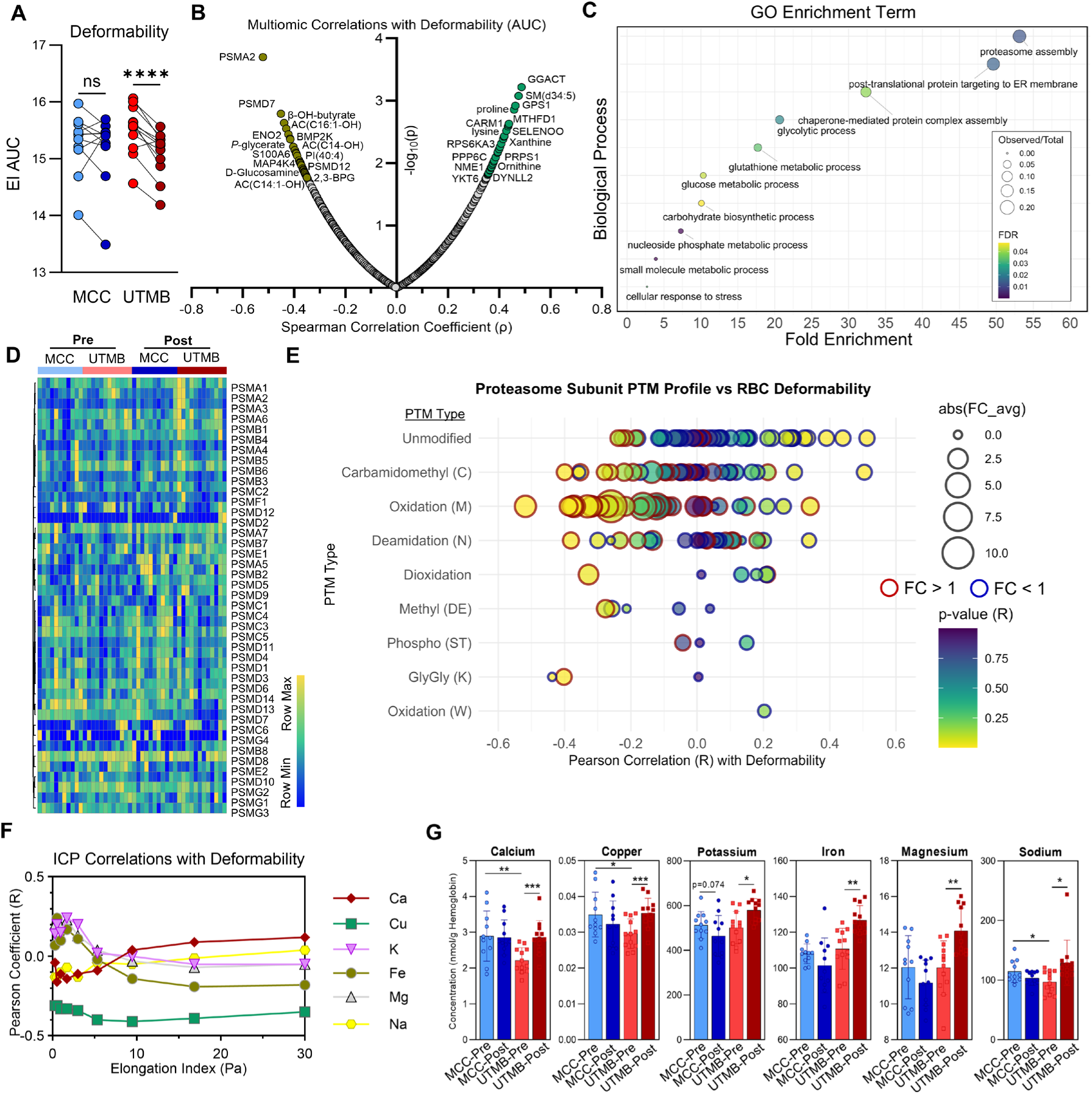
Molecular Correlates with RBC Deformability. A) Total deformability decreases specifically after UTMB (see Sup Fig for deform on all Pressures). B) Spearman correlations with Deformability AUC. C) GO Enrichment terms with deformability. D) Heatmap showing total levels of proteasome subunits. (E) Dot plot showing the Pearson correlation with RBC deformability on the x-axis. The size of each dot is proportional to the Post/Pre fold change and the dot outline indicates with the fold change increased (red) or decreased (blue) after the race. The significance of the correlation is indicated by color. (F) Pearson correlation at each shear stress pressure for metals, along with (G) abundance before and after each race.

### Static abundant plasma analytes and increasing free bilirubin indicate increased extravascular hemolysis during ultra-running

In consideration of the documented expansion of plasma fluid volumes during ultrarunning (*44*), we re-processed plasma proteomics data without cross run normalization (Supplementary Table – a processing step which standardizes total protein signal in each sample across the analyzed dataset (*45*). This step was employed to analyze protein trends with respect to possible changes in total proteome content during the MCC and UTMB events. Non-normalized plasma proteomics data was thus mined for markers of hemolysis or blood volume increase to further interrogate the decreases in hematocrit observed specifically after the UTMB (**Figure 3A**). Despite static levels of cell free hemoglobin (HBA and HBB), intravascular hemolysis may have occurred in light of decreased haptoglobin (HP) levels in both races, and decreased transferrin (TF) after the MCC (**Figure 8A**), the effects of which have been documented in distance runners (*10*). The most abundant plasma proteins in the proteomics dataset including albumin and apolipoprotein A1 (ApoA-I) were unchanged in either race **(Figure 8A**). In addition, plasma levels of sodium, iron, and calcium measured by ICP-MS were unchanged (**Figure 8B**). To identify molecules that associate with Pre and Post hematocrit, a linear mixed model was prepared that corrected for age, sex, and BMI (**Figure 8C**). The top associated feature with hematocrit was the ketone body acetoacetate. In addition, both plasma bilirubin (total) and hypoxanthine emerged as associated features after correction. Plasma bilirubin is a marker of extravascular hemolysis (*46, 47*), and was a significant inverse correlate with hematocrit along with plasma hypoxanthine (**Figure 8D**). Hypoxanthine also increased in plasma after the UTMB (**Figure 8D**), mirroring RBC alterations (**Figure 5A**). In addition, RBC creatinine, a marker of RBC circulatory age (*48, 49*), was elevated in RBCs after both races, though to a significantly higher extent after the MCC (**Figure 8E**).

**Figure 8.**
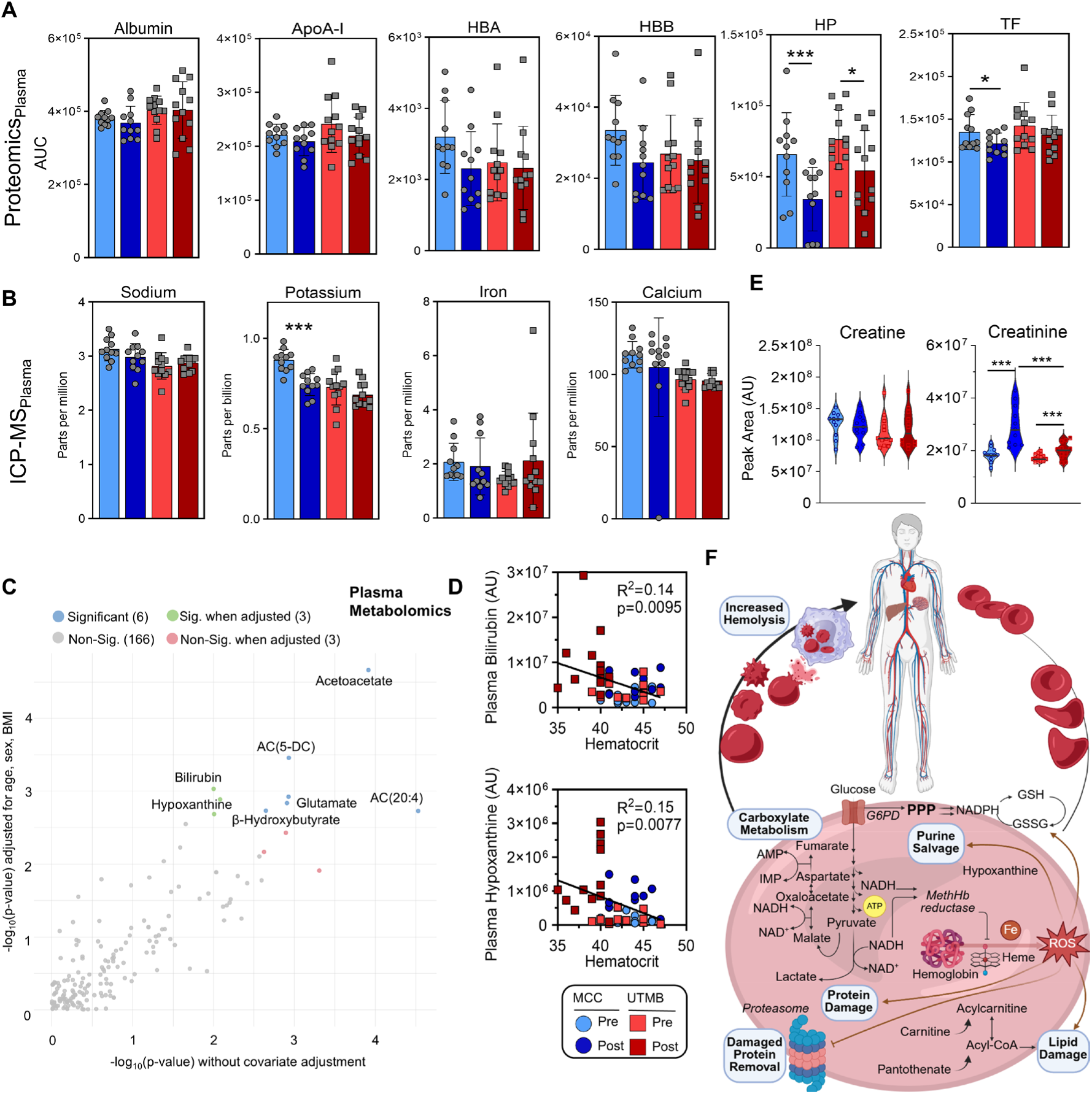
Multi-Omics signatures in plasma highlight increased hemolysis after ultrarunning. (A) The plasma levels of abundant proteins albumin, apolipoprotein A1 (ApoA-I), hemoglobin alpha (HBA) and beta (HBB), haptoglobin (HP), and transferrin (TF). (B) Plasma levels of trace elements. (C) Linear mixed model of hematocrit corrected for age, sex, height, and weight using plasma metabolomics data. (D) Pearson correlation plots of hematocrit and plasma levels of bilirubin and hypoxanthine. Individual races and time points are indicated. (E) RBC levels of creatine and creatinine. (F) A proposed model of RBC damage accumulation during long distance trail running.

## Discussion

Here we describe the impact of marathon and ultramarathon trail running distances on multi-omic (metabolomic, lipidomic, proteomic, and metallomic) profiles in plasma and RBC compartments, and how these profiles correspond to alterations in hemorheological parameters. Our findings detail the effects of ultra-running on a cascade of molecular events that extend beyond mechanical trauma, implicating systemic inflammation, oxidative stress, and metabolic remodeling as key drivers of RBC aging and dysfunction. Signatures implicate a mounting immune response that begins during a 40 km run (MCC) and accumulates during a 171 km ultramarathon (UTMB). This inflammation manifests across multiple measurement modalities, from an increase in total white blood cell count, to elicitation of a cytokine response involving IL-6, to stimulation of the acute phase response, to production of proinflammatory metabolites such as kynurenine. These molecular signatures correspond with an initial increase after the MCC in externalized phosphatidylserine (PS) on the RBC membrane, which is a marker for RBC clearance from circulation (*50*). While elevations of external PS were not apparent in circulating RBCs after the longer UTMB race, these cells were less deformable, accompanied by decreased cell volume, higher oxidation of structural proteins, increased generation of microparticles and release of immature reticulocytes. Perhaps the most observable difference between the MCC and UTMB was decreased hematocrit during the latter race. While some studies have suggested RBC are removed substantially due ultrarunning (*51, 52*), use of more direct carbon-monoxide rebreathing techniques that assess total hemoglobin mass and total red blood cell volume after ultrarunning (including the UTMB) have attributed hematocrit alterations to plasma volume expansion of 18-20% (*44, 53*). Therefore, hemodilution is likely a large contributor to the decreased hematocrit observed immediately following the UTMB runners studied herein. Rather, these results are consistent with a model positing acceleration of damage to RBC during running that results in increased splenic sequestration and extravascular hemolysis of less deformable, more damaged RBCs, in keeping with similar findings in the context of transfusion of RBCs after prolonged storage (*41*). Consistent with this hypothesis, plasma levels of bilirubin increased in accordance with decreased hematocrit, and while cell free hemoglobin remained unchanged, haptoglobin was decreased. As RBCs cross a threshold of accumulated oxidative and mechanical damage, they are selected for removal from circulation in the liver or spleen via macrophage phagocytosis (*54*). During periods of excess RBC removal, indirect bilirubin is released back into circulation rather than removed through the biliary duct and intestine (*46*). Therefore, these results illustrate how extended endurance running exercise results in an accelerated aging process within the RBC population, thereby leading to enhanced removal of damaged cells from circulation.

Running-induced RBC damage can stem from increased shear stress in the foot leading to mechanical trauma (*55*). Several non-mechanical injuries to RBC during exercise have also been documented that impair RBC membrane structure and deformability while promoting fragility. These mechanisms include alteration of the membrane lipidome (*7, 56*), cytoskeleton (*52, 57*), and depletion of antioxidant systems (*58*), which occur in conjunction with increased osmotic fragility due, in part, to prolonged exposure to lactate acidosis and subsequent dehydration at higher exercise intensities (*59–63*). Indeed, RBCs here decreased in volume and deformability, and while glutathione did not appear to be depleted, protein and lipid oxidation increased substantially.

We observed multiple impacts of running on the RBCs membrane lipidome. RBCs repair membrane lipid damage using the Lands Cycle, which relies on the generation of acyl-CoA substrates to replace damaged acyl chains (*64*). This process is buffered by acylcarnitines, which serve as an ATP-independent source of acyl chains (*65–67*). Therefore, carnitine availability can bolster this process and improve RBC function under extreme oxidative stress environments of blood storage (*68*). In this study, however, we observed carnitine accumulation specifically after UTMB, which might suggest that RBCs are utilizing the acyl-CoA pool at an accelerated rate and of the CoA precursor pantothenate (Vitamin B5), which is metabolized by RBCs (*69, 70*), suggests alternative strategies to maintain the Lands Cycle during exercise provided that intestinal absorption is optimized (*71, 72*).

Double bonds in acyl chains are susceptible to hydroxylation via alpha-oxidation to generate hydroxyacylcarnitines. Inverse correlations between hydroxyacylcarnitines and RBC deformability seen here were also observed in cyclists (*7*), who do not experience similar mechanical stress in the foot, thereby indicating alternative sources of oxidative lipid stress. Notably, these molecules increase in RBC as a function of cardiorespiratory workload in a controlled cardiopulmonary exercise test (CPET) in healthy subjects (*73*). Acylcarnitine hydroxylation may occur in conjunction with increased desaturation of fatty acids, which are subsequently susceptible to lipid peroxidation (*74*). Indeed, fatty acid desaturase activity is upregulated in part to recycle NAD^+^ (*75*). While the role of this pathway has been established in *ex vivo* RBCs storage (*34*), future work should look at the role this pathway plays in response to exercise *in vivo*.

The extent of oxidation on proteasome subunits inversely correlated with RBC elongation index, suggesting that the loss of ability to degrade damaged proteins may be linked to loss of deformability, or that deformability itself may influence proteasome activity. Indeed, loss of proteasome activity in conjunction with increasing oxidative stress and decreasing RBC deformability has been observed in ultra-trail running races on La Reunion Island. (*76*) In addition, the levels of copper were consistently inverse correlated with elongation index at all shear stresses. Copper accumulation is known to occur as part of the acute phase response (*77*) and can directly decrease RBC deformability (*78*) through interaction with and subsequent oxidation of membrane phospholipids (*79*). Future work should focus on understanding the source of this copper (i.e. extracellular versus loss of conjugation to copper-containing proteins such as superoxide dismutase, for example), its direct role in RBC function, and how exercise or other stressors influence concentration dynamics.

These results, however, do highlight the importance of oxidative stress management in circulation prior to RBC removal by the reticuloendothelial system. One process by which RBCs are selected for removal involves their ability to traverse the interendothelial slits within the spleen. With sub-1.5 µm diameters, this transit is thus invariably dependent on the RBCs ability to deform (*80*). Moreover, passing through gradients of varying tonicity in both the spleen and kidney requires that the RBCs can maintain resilience to osmotic stress. Interestingly, RBCs that possess high levels of kynurenine are also increasingly fragile to changes in osmotic gradients (*28*). Because kynurenine is not synthesized by RBCs, its abundance is therefore a reflection of the circulating environment. Individuals who donate RBC units with higher levels of kynurenine tend to be of older age or higher BMI, and the plasma proteome within these units is characterized by a sterile inflammatory environment (*28*). This signature is also present in interferon-associated inflammation of Trisomy 21 (*81*). In an acutely inflamed environment such as COVID-19, kynurenine is strongly correlated with both IL-6 levels and disease severity (*25, 27, 82*). Notably, comparable damage to RBC membrane lipids and proteins has been documented in COVID-19 patients (*83*), in line with decreased RBC deformability (*84, 85*). These studies, however, were performed using blood from one or two collections and thus represented “snapshots” of individual biology at the time of collection. In this study, we were able to obtain a more dynamic view of this network after paired sampling in the same subjects. Collectively, the association between cell rheology and oxidative damage to proteins and lipids reinforces previous findings, though expands these from therapeutic settings such as blood storage and pathological settings such as COVID-19 to a new setting of prolonged endurance exercise.

The present findings not only confirm that extreme endurance exercise induces a complex cascade of RBC damage and repair processes but also offer mechanistic insights that have broader implications. Repeated exposure to mechanical trauma (e.g., foot-strike hemolysis) and additional that is evident in the accumulation of oxidized proteins (e.g., methionine oxidation) and altered lipid profiles. In addition to these mechanistic insights, our study underscores important differences between running-based endurance activities and other modalities such as cycling. Running, particularly on hard surfaces and in challenging trail environments, uniquely exacerbates mechanical damage to RBCs through repetitive impact forces. In contrast, non–weight-bearing activities predominantly induce shear stress without the added burden of direct mechanical trauma. This distinction is critical when considering the design of training programs and recovery strategies, as athletes engaged in running sports may require targeted interventions—such as optimized nutritional support (e.g., antioxidants, pantothenate, or carnitine supplementation) or tailored recovery protocols—to minimize RBC damage and maintain optimal oxygen transport capacity.

Moreover, the multi-omics signatures identified here have direct implications for athletic performance and recovery. The observed decreases in RBC deformability and alterations in metabolite and protein profiles suggest that prolonged endurance exercise challenges the balance between RBC damage and repair. In the short term, this imbalance may impair microcirculatory flow and reduce oxygen delivery, contributing to performance decline during ultramarathons. Over the longer term, repeated bouts of exercise-induced RBC turnover could drive hematological adaptations that, while beneficial (e.g., a younger, more resilient RBC pool), also risk cumulative damage if recovery is insufficient. These insights pave the way for future research aimed at defining an “optimal range” of hemolysis and RBC turnover during exercise—a concept that could be instrumental in developing personalized training and nutritional interventions.

Inter-individual variability, driven by factors such as age, sex, training status, and genetic predisposition likely influences the degree of molecular damage observed, highlighting the need for personalized approaches in exercise regimens and recovery strategies. Moreover, the parallels between the molecular signatures observed in endurance athletes and those in other models of oxidative stress—such as sepsis, aging, or even storage lesions in transfused blood—offer broader biological insights that could extend to clinical settings, including transfusion medicine. Finally, while the current study provides a comprehensive snapshot of the acute molecular responses to long-distance running, future longitudinal studies are needed to assess the persistence of these changes and to directly link molecular alterations to functional outcomes. Such work would not only deepen our understanding of exercise-induced cellular stress but also inform interventions aimed at preserving RBCs functionality and overall cardiovascular health in athletes.

## Materials and Methods

### Sample Collection, Hematological, and Hemorheological Measurements

Sample collection and analyses have been previously described (*14*). Briefly, blood samples were collected from endurance-trained participants before and after either a 40 km trail race (MCC) or a 171 km ultra-trail (UTMB), under similar environmental conditions. The protocol was approved by the Ethics Committee (CPP Ouest VI, ethics committee agreement 19.03.14.41740 received on 05/02/2019), and the study was conducted in accordance with the Declaration of Helsinki. Standardized protocols were used to assess hematological, hemorheological, and red blood cell senescence parameters, including blood viscosity, RBC deformability and aggregation, plasma free hemoglobin, microparticle release, phosphatidylserine exposure, and inflammatory markers such as IL-6. Full methodological details have been described previously

### Metabolomics and Lipidomics Sample Preparation

Prior to metabolomics or lipidomics analysis, individual samples were placed in 1.5 ml tubes for either metabolite or lipid extraction. Twenty µL were suspended in 180 µL of water/methanol (50:50 v/v) for metabolite extraction or 180 µL of isopropanol/methanol (50:50 v/v) for lipid extraction. Suspensions were vortexed for 30 minutes at 4°C and then centrifuged for 10 minutes,

### Metabolomics and Lipidomics UHPLC-MS data acquisition and processing

Analyses were performed as previously published. (*86, 87*) Briefly, the analytical platform employs a Vanquish UHPLC system (Thermo Fisher Scientific, San Jose, CA, USA) coupled online to a Q Exactive mass spectrometer (Thermo Fisher Scientific, San Jose, CA, USA). Polar metabolite extracts are resolved in singlicate over a Kinetex C18 column, 2.1 x 150 mm, 1.7 µm particle size (Phenomenex, Torrance, CA, USA) equipped with a guard column (SecurityGuard^TM^ Ultracartridge – UHPLC C18 for 2.1 mm ID Columns, Phenomenex, Torrance, CA, USA) using an aqueous phase (A) of water and 0.1% formic acid and a mobile phase (B) of acetonitrile and 0.1% formic acid for positive ion polarity mode, and an aqueous phase (A) of water:acetonitrile (95:5) with 1 mM ammonium acetate and a mobile phase (B) of acetonitrile:water (95:5) with 1 mM ammonium acetate for negative ion polarity mode. The column is equilibrated at 5% B, and upon injection of 10 μl of ex-tract, samples are eluted from the column using the solvent gradient: 0.5-1.1 min 5-95% B at 0.45 mL/min; hold at 95% B for 1.65 min at 0.45 mL/min, and then decrease to 5% over 0.25 min at 0.45 ml/min, followed by a re-equilibration hold at 5% B for 2 minutes at 0.45 ml/min. The Q Exactive mass spectrometer (Thermo Fisher Scientific, San Jose, CA, USA) is operated independently in positive or negative ion mode, scanning in Full MS mode (2 μscans) from 60 to 900 m/z at 70,000 resolution, with 4 kV spray voltage, 45 sheath gas, 15 auxiliary gas, AGC target = 3e6, maximum IT = 200 ms. Non-polar lipid extracts are re-solved over an ACQUITY HSS T3 column (2.1 x 150 mm, 1.8 µm particle size (Waters, MA, USA) using an aqueous phase (A) of 25% acetonitrile and 5 mM ammonium acetate and a mobile phase (B) of 90% isopropanol, 10% acetonitrile and 5 mM ammonium acetate. The column is equilibrated at 30% B, and upon injection of 10 μl of extract, samples are eluted from the column using the solvent gradient: 0-9 min 30-100% B at 0.325 mL/min; hold at 100% B for 3 min at 0.3 mL/min, and then decrease to 30% over 0.5 min at 0.4 ml/min, followed by a re-equilibration hold at 30% B for 2.5 minutes at 0.4 ml/min. The Q Exactive mass spectrometer (Thermo Fisher) is operated in positive ion mode, scanning in Full MS mode (2 μscans) from 150 to 1500 m/z at 70,000 resolution, with 4 kV spray voltage, 45 sheath gas, 15 auxiliary gas. When required, dd-MS2 is performed at 17,500 resolution, AGC target = 1e5, maximum IT = 50 ms, and stepped NCE of 25, 35 for positive mode, and 20, 24, and 28 for negative mode. Calibration is performed prior to analysis using the Pierce^TM^ Positive and Negative Ion Calibration Solutions (Thermo Fisher Scientific).

### Metabolomics and Lipidomics Data analysis

Acquired data were converted from raw to mzXML file format using Mass Matrix (Cleveland, OH, USA). Samples were analyzed in randomized order with a technical mixture injected after every 10 samples to qualify instrument performance. Metabolite assignments are performed using accurate intact mass (sub-10 ppm), Isotopologue distributions, and retention time/spectral comparison to an in-house standard compound library (MSMLS, IROA Technologies, NJ, USA) using MAVEN (Princeton, NJ, USA). Lipidomics data are analyzed using LipidSearch 5.0 (Thermo Scientific), which provides lipid identification on the basis of accurate intact mass, isotopic pattern, and fragmentation pattern to determine lipid class and acyl chain composition. Graphs, heat maps and statistical analyses (either T-Test or ANOVA), multivariate analyses including Principal Component Analysis (PCA), Partial Least Squares-Discriminant Analysis (PLS-DA), hierarchical clustering analysis (HCA), and metabolite pathway enrichment analysis are performed using MetaboAnalyst 5.0 (*88*) DSPC IL-6 network filtered by degree (1.5), betweenness (1.0), and correlation (p < 0.001) were plotted using CytoScape. (*89*)

### Inductively- Coupled Plasma (ICP) Mass Spectrometry

Samples were analyzed as previously published. (*90*) Prior to ICP-MS analysis, 10 μL of plasma were aliquoted into a 15 mL conical tube. 200 μL of 65% nitric acid and 20 ng/mL of gold were added into each sample followed by and addition of 100 μL of 30% hydrogen peroxide and brief vortexing. Samples were then incubated in an oven at 70°C for approximately 2 h. Following incubation, 2190 μL of MilliQ water was added to each tube (final nitric acid percentage of ∼5%) and all samples were vortexed briefly. All sam-ples were then diluted 1:15 in a solution consisting of 20 ng/mL and 5% nitric acid. Final dilutions of 1:250 and 1:3750 were then analyzed via ICP-MS. Different dilutions were used to ensure all analytes fell within the calibration curves. All chemicals and materials used for ICP-MS analysis were obtained from Thermo Fisher and all ICP-MS calibrants and solutions were obtained from SPEX CertiPrep. All samples were analyzed on a Thermo Scientific iCAP RQ ICP-MS coupled to a ESI SC-4DX FAST autosampler system utilizing a peristaltic pump. The optimization of the system was performed before the run by first calibrating the system with ICP-MS iCAP Q/Qnova Calibration Solution, Specpure. The system was subsequently tuned using a tuning solution consisting of Ba, Bi, Ce, Co, In, Li, and U at 1.00 ± 0.05 μg/L. To monitor performance while the system was running, we continually pumped internal standard mix via the peristaltic pump and monitored signal throughout the run. Additionally, quality controls of a known concentration of each analyte were injected at the beginning, throughout the run between samples, and at the end of the run. Acceptance criteria for all QCs was ±25% of the known concentration. Thermo Scientific Qtegra software was used for all data acquisition and analysis.

### Proteomics Sample Preparation

Plasma and RBC samples were digested using S-Trap 96-well plates (Protifi, Huntington, NY) following the manufacturer’s protocol. Briefly, approximately 50 µg of protein was mixed with 5% SDS. Samples were reduced with 10 mM dithiothreitol (DTT) at 55°C for 30 minutes, cooled to room temperature, and alkylated with 25 mM iodoacetamide (IAM) in the dark for 30 minutes.Next, phosphoric acid was added to achieve a final concentration of 2.5 % followed by six volumes of binding buffer (90% methanol, 100 mM triethylammonium bicarbonate [TEAB], pH 7.1). After gentle mixing, the protein solution was loaded onto an S-Trap 96-well plate and centrifuged at 1500 × g for 2 minutes. The plate was then washed three times with 300 µL of binding buffer. For digestion, 1 µg of sequencing-grade trypsin (Promega) and 125 µL of digestion buffer (50 mM TEAB) were added to the filter, and samples were incubated at 37°C for 6 hours. Peptides were sequentially eluted using 100 µL of each of the following buffers: (1) 50 mM TEAB, (2) 0.2% formic acid in water, and (3) 50% acetonitrile with 0.2% formic acid in water. The peptide solutions were pooled, lyophilized, and resuspended in 500 µL of 0.1% formic acid (FA)

### Proteomics Data acquisition and processing

A 20 µL aliquot of each sample was loaded onto individual Evotips for desalting, followed by three washes with 200 µL of 0.1% formic acid (FA). To maintain hydration until analysis, 100 µL of storage solvent (0.1% FA) was added to each Evotip. Peptide separation was performed using a PepSep column (150 µm inner diameter, 15 cm) packed with ReproSil C18 (1.9 µm, 120Å resin) and a pre-set 30 samples-per-day gradient on the Evosep One system (Evosep, Odense, Denmark). The Evosep One system was coupled to a timsTOF Pro mass spectrometer (Bruker Daltonics, Bremen, Germany) via a CaptiveSpray nano-electrospray ion source (Bruker Daltonics). The mass spectrometer was operated in DIA-PASEF mode. We used a method with four windows in each 100 ms dia-PASEF scan. 32 of these scans covered the diagonal scan line for doubly and triply charged peptides in the m/z - ion mobility plane with narrow 25 m/z precursor windows.

A project specific spectral library generated from 24 high-pH reverse-phase peptide fractions acquired with DDA-PASEF acquisition. MS data were collected over an m/z range of 100 to 1,700. During each MS/MS data collection, each TIMS cycle was 1.17 seconds and included 1 MS and 10 PASEF MS/MS scans. Low-abundance precursor ions with an intensity above a threshold of 500 counts but below a target value of 20000 counts were repeatedly scheduled and otherwise dynamically excluded for 0.4 min. Pooled plasma or RBCs digest was separated by high pH reversed phase chromatography on a Gemini-NH C18, 4.6 x 250 mm analytical column containing 3 μM particles, flow rate was 0.6 mL/min. The solvent consisted of 20 mM ammonium bicarbonate (pH 10) as mobile phase (A) and 20 mM ammonium bicarbonate and 75% ACN (pH 10) as mobile-phase B. Sample separation was accomplished using the following linear gradient: from 0 to 5% B in 10 min, from 5 to 50% B in 80 min, from 50 to 100% B in 10 min, and held at 100% B for an additional 10 min. 96 fractions were collected along with the LC separation and were concatenated into 24 fractions by combining fractions 1, 25, 49, 73 and so on. Samples were dried in Speed-Vac, resuspended in 80 ul of 0.1% FA and 20ul of each fraction was loaded onto individual Evotips.

### Proteomics Data Processing

Raw .d files were analyzed with Spectronaut 17.6 (Biognosys) using default settings. For plasma analyses taking into consideration plasma volume expansion, searches were performed using identical settings except for de-selecting Cross Run Normalization. Spectral libraries were prepared searching for carbamidomethyl (C), Glu>PyroGlu, and Acetyl (N-ter) as fixed modifications, with variable modifications including methyl (D,E), deamidation (N), oxidation (M), His>Asp, Phospho (ST), oxidation (W), dioxidation, and GlyGly (K). Proteomics GO networks were prepared using ShinyGO 0.82 (*91, 92*)

## Author contributions

Conceptualization: PC, AD Methodology: TN, FC, DS, MD

Investigation: TN, ES, FC, EN, MR, SS, PR, GM, PC, DS, MD

Visualization: TN

Supervision: PC, AD, TN, KCH

Writing—original draft: TN, AD

Writing—review & editing: All authors

## Data and materials availability

All data are available in the main text or the supplementary materials.

## Supplementary Materials

Supplementary materials are provided as 5 Tables (spreadsheet xlsx format) and 7 figures (pdf).

## References

1. H. Mairbäurl, Red blood cells in sports: effects of exercise and training on oxygen supply by red blood cells. Frontiers in Physiology 4, 332 (2013).

2. T. Yoshida, M. Prudent, A. D’Alessandro, Red blood cell storage lesion: causes and potential clinical consequences. Blood Transfusion, (2019).

3. A. D’Alessandro, K. C. Hansen, E. Z. Eisenmesser, J. C. Zimring, Protect, repair, destroy or sacrifice: a role of oxidative stress biology in inter-donor variability of blood storage? Blood Transfus 17, 281–288 (2019).

4. M. Hargreaves, L. L. Spriet, Skeletal muscle energy metabolism during exercise. Nat Metab 2, 817–828 (2020).

5. S. Amatori et al., Are pre-race serum blood biomarkers associated with the 24-h ultramarathon race performance? European Journal of Sport Science 24, 431–439 (2024).

6. A. Siekierzycka et al., Plasma cardiovascular stress biomarkers response to marathon running. Sports Medicine and Health Science, (2024).

7. T. Nemkov et al., Acute Cycling Exercise Induces Changes in Red Blood Cell Deformability and Membrane Lipid Remodeling. Int J Mol Sci 22, (2021).

8. I. San-Millán et al., Metabolomics of Endurance Capacity in World Tour Professional Cyclists. Frontiers in Physiology 11, 578 (2020).

9. R. Fleischer, Ueber eine neue form von haemoglobinurie beim menschen. Berl Klin Wschr 18, 691 (1881).

10. A. Gartner, N. Dombrowski, N. Lowe, V. Behzadpour, R. Zackula, Foot-strike Hemolysis: A Scoping Review of Long-Distance Runners. Kans J Med 17, 119–124 (2024).

11. D. Nash et al., IL-6 signaling in acute exercise and chronic training: Potential consequences for health and athletic performance. Scand J Med Sci Sports 33, 4–19 (2023).

12. H. Northoff, A. Berg, Immunologic mediators as parameters of the reaction to strenuous exercise. Int J Sports Med 12 Suppl 1, S9–15 (1991).

13. E. Nader et al., Impact of a 10 km running trial on eryptosis, red blood cell rheology, and electrophysiology in endurance trained athletes: a pilot study. European Journal of Applied Physiology 120, 255–266 (2020).

14. M. Robert et al., Impact of Trail Running Races on Blood Viscosity and Its Determinants: Effects of Distance. Int J Mol Sci 21, (2020).

15. S. Skinner et al., Differential impacts of trail and ultra-trail running on cytokine profiles: An observational study. Clin Hemorheol Microcirc 78, 301–310 (2021).

16. K. Contrepois et al., Molecular Choreography of Acute Exercise. Cell 181, 1112–1130.e1116 (2020).

17. C. C. F. Howe et al., Untargeted Metabolomics Profiling of an 80.5 km Simulated Treadmill Ultramarathon. Metabolites 8, (2018).

18. Z. Stander et al., The unaided recovery of marathon-induced serum metabolome alterations. Sci Rep 10, 11060 (2020).

19. T. B. Høeg, K. Chmiel, A. E. Warrick, S. L. Taylor, R. H. Weiss, Ultramarathon Plasma Metabolomics: Phosphatidylcholine Levels Associated with Running Performance. Sports (Basel) 8, (2020).

20. H. Mairbäurl, Red blood cells in sports: Effects of exercise and training on oxygen supply by red blood cells. Frontiers in Physiology 4, (2013).

21. C. Gaud et al., BioPAN: a web-based tool to explore mammalian lipidome metabolic pathways on LIPID MAPS. F1000Res 10, 4 (2021).

22. Y. Chen, J. Zhang, W. Cui, R. L. Silverstein, CD36, a signaling receptor and fatty acid transporter that regulates immune cell metabolism and fate. J Exp Med 219, (2022).

23. T. G. Manfredi et al., Plasma creatine kinase activity and exercise-induced muscle damage in older men. Med Sci Sports Exerc 23, 1028–1034 (1991).

24. S. Basu et al., Sparse network modeling and metscape-based visualization methods for the analysis of large-scale metabolomics data. Bioinformatics 33, 1545–1553 (2017).

25. D. A. Bizjak et al., Kynurenine serves as useful biomarker in acute, Long- and Post-COVID-19 diagnostics. Front Immunol 13, 1004545 (2022).

26. J. P. Dewulf et al., Urine metabolomics links dysregulation of the tryptophan-kynurenine pathway to inflammation and severity of COVID-19. Sci Rep 12, 9959 (2022).

27. T. Thomas, et al., COVID-19 infection alters kynurenine and fatty acid metabolism, correlating with IL-6 levels and renal status. JCI Insight 5, (2020).

28. T. Nemkov et al., Regulation of kynurenine metabolism by blood donor genetics and biology impacts red cell hemolysis in vitro and in vivo. Blood 143, 456–472 (2024).

29. E. Nader et al., Eryptosis and hemorheological responses to maximal exercise in athletes: Comparison between running and cycling. Scandinavian Journal of Medicine & Science in Sports 28, 1532–1540 (2018).

30. E. Lang, F. Lang, Triggers, Inhibitors, Mechanisms, and Significance of Eryptosis: The Suicidal Erythrocyte Death. BioMed Research International 2015, 1–16 (2015).

31. T. Nemkov, S. M. Qadri, W. P. Sheffield, A. D’Alessandro, Decoding the metabolic landscape of pathophysiological stress-induced cell death in anucleate red blood cells. Blood Transfusion, (2020).

32. A. D’Alessandro et al., Ferroptosis regulates hemolysis in stored murine and human red blood cells. Blood 145, 765–783 (2025).

33. M. A. Perez et al., Ether lipid deficiency disrupts lipid homeostasis leading to ferroptosis sensitivity. PLoS Genet 18, e1010436 (2022).

34. T. Thomas et al., Fatty acid desaturase activity in mature red blood cells and implications for blood storage quality. Transfusion, (2021).

35. A. D’Alessandro et al., Functional and multi-omics signatures of mitapivat efficacy upon activation of pyruvate kinase in red blood cells from patients with sickle cell disease. Haematologica 109, 2639–2652 (2024).

36. T. Nemkov et al., Hypoxia modulates the purine salvage pathway and decreases red blood cell and supernatant levels of hypoxanthine during refrigerated storage. Haematologica 103, 361–372 (2018).

37. C. Chen et al., Erythrocyte ENT1-AMPD3 Axis is an Essential Purinergic Hypoxia Sensor and Energy Regulator Combating CKD in a Mouse Model. J Am Soc Nephrol 34, 1647–1671 (2023).

38. T. Nemkov et al., Metabolism of Citrate and Other Carboxylic Acids in Erythrocytes As a Function of Oxygen Saturation and Refrigerated Storage. Frontiers in Medicine 4, 175 (2017).

39. A. D’Alessandro et al., Donor sex, age and ethnicity impact stored red blood cell antioxidant metabolism through mechanisms in part explained by glucose 6-phosphate dehydrogenase levels and activity. Haematologica, haematol.2020.246603 (2020).

40. R. O. Francis et al., Donor glucose-6-phosphate dehydrogenase deficiency decreases blood quality for transfusion. J Clin Invest 130, 2270–2285 (2020).

41. S. Peltier et al., Proteostasis and metabolic dysfunction characterize a subset of storage-induced senescent erythrocytes targeted for post-transfusion clearance. J Clin Invest, (2025).

42. J. A. Reisz et al., Methylation of protein aspartates and deamidated asparagines as a function of blood bank storage and oxidative stress in human red blood cells. Transfusion 58, 2978–2991 (2018).

43. Z. Lv et al., Domain alternation and active site remodeling are conserved structural features of ubiquitin E1. J Biol Chem 292, 12089–12099 (2017).

44. P. Robach et al., Hemolysis induced by an extreme mountain ultra-marathon is not associated with a decrease in total red blood cell volume. Scand J Med Sci Sports 24, 18–27 (2014).

45. R. Bruderer et al., Optimization of Experimental Parameters in Data-Independent Mass Spectrometry Significantly Increases Depth and Reproducibility of Results. Mol Cell Proteomics 16, 2296–2309 (2017).

46. F. Rapido et al., Prolonged red cell storage before transfusion increases extravascular hemolysis. J Clin Invest 127, 375–382 (2017).

47. G. Lippi, M. Plebani, S. Di Somma, G. Cervellin, Hemolyzed specimens: a major challenge for emergency departments and clinical laboratories. Crit Rev Clin Lab Sci 48, 143–153 (2011).

48. I. Syllm-Rapoport, A. Daniel, H. Starck, A. Hartwig, J. Gross, Creatine in density-fractionated red cells, a useful indicator of erythropoietic dynamics and of hypoxia past and present. Acta Haematol 66, 86–95 (1981).

49. C. P. Ku, H. Passow, Creatine and creatinine transport in old and young human red blood cells. Biochim Biophys Acta 600, 212–227 (1980).

50. F. E. Boas, L. Forman, E. Beutler, Phosphatidylserine exposure and red cell viability in red cell aging and in hemolytic anemia. Proc Natl Acad Sci U S A 95, 3077–3081 (1998).

51. W. H. Reinhart, M. Staubli, P. W. Straub, Impaired red cell filterability with elimination of old red blood cells during a 100-km race. Journal of Applied Physiology 54, 827–830 (1983).

52. A. Yusof et al., Exercise-induced hemolysis is caused by protein modification and most evident during the early phase of an ultraendurance race. J Appl Physiol (1985) 102, 582–586 (2007).

53. B. Krumm et al., Accelerated Red Blood Cell Turnover Following Extreme Mountain Ultramarathon? Med Sci Sports Exerc 57, 904–911 (2025).

54. T. R. Klei, S. M. Meinderts, T. K. van den Berg, R. van Bruggen, From the Cradle to the Grave: The Role of Macrophages in Erythropoiesis and Erythrophagocytosis. Front Immunol 8, 73 (2017).

55. R. D. Telford et al., Footstrike is the major cause of hemolysis during running. J Appl Physiol (1985) 94, 38–42 (2003).

56. T. Yamada, M. Tohori, T. Ashida, N. Kajiwara, H. Yoshimura, Comparison of effects of vegetable protein diet and animal protein diet on the initiation of anemia during vigorous physical training (sports anemia) in dogs and rats. J Nutr Sci Vitaminol (Tokyo) 33, 129–149 (1987).

57. R. Beneke, D. Bihn, M. Hütler, R. M. Leithäuser, Haemolysis caused by alterations of alpha- and beta-spectrin after 10 to 35 min of severe exercise. Eur J Appl Physiol 95, 307–312 (2005).

58. J. A. Smith, Exercise, Training and Red Blood Cell Turnover:. Sports Medicine 19, 9–31 (1995).

59. O. S. Platt, S. E. Lux, D. G. Nathan, Exercise-induced hemolysis in xerocytosis. Erythrocyte dehydration and shear sensitivity. J Clin Invest 68, 631–638 (1981).

60. W. van Beaumont, S. Underkofler, S. van Beaumont, Erythrocyte volume, plasma volume, and acid-base changes in exercise and heat dehydration. Journal of Applied Physiology 50, 1255–1262 (1981).

61. V. Lipovac, M. Gavella, Z. Turk, Z. Škrabalo, Influence of lactate on the insulin action on red blood cell filterability. Clinical Hemorheology and Microcirculation 5, 421–428

62. J. A. Smith, R. D. Telford, M. Kolbuch-Braddon, M. J. Weidemann, Lactate/H+ uptake by red blood cells during exercise alters their physical properties. European Journal of Applied Physiology and Occupational Physiology 75, 54–61 (1997).

63. P. Connes et al., Opposite effects of in vitro lactate on erythrocyte deformability in athletes and untrained subjects. Clinical Hemorheology and Microcirculation 31, 311–318 (2004).

64. W. E. Lands, Metabolism of glycerolipides; a comparison of lecithin and triglyceride synthesis. The Journal of Biological Chemistry 231, 883–888 (1958).

65. A. Arduini et al., Role of carnitine and carnitine palmitoyltransferase as integral components of the pathway for membrane phospholipid fatty acid turnover in intact human erythrocytes. The Journal of Biological Chemistry 267, 12673–12681 (1992).

66. A. Arduini et al., Effect of L-carnitine and acetyl-L-carnitine on the human erythrocyte membrane stability and deformability. Life Sciences 47, 2395–2400 (1990).

67. A. Arduini et al., Acyl-trafficking in membrane phospholipid fatty acid turnover: The transfer of fatty acid from the acyl-L-carnitine pool to membrane phospholipids in intact human erythrocytes. Biochemical and Biophysical Research Communications 187, 353–358 (1992).

68. T. Nemkov et al., Genetic regulation of carnitine metabolism controls lipid damage repair and aging RBC hemolysis in vivo and in vitro. Blood, (2024).

69. K. F. Annous, W. O. Song, Pantothenic acid uptake and metabolism by red blood cells of rats. The Journal of Nutrition 125, 2586–2593 (1995).

70. S. A. Mian et al., Vitamin B5 and succinyl-CoA improve ineffective erythropoiesis in SF3B1-mutated myelodysplasia. Sci Transl Med 15, eabn5135 (2023).

71. M. H. Williams, Vitamin supplementation and athletic performance. International Journal for Vitamin and Nutrition Research. Supplement = Internationale Zeitschrift Fur Vitamin- Und Ernahrungsforschung. Supplement 30, 163–191 (1989).

72. M. J. Webster, Physiological and performance responses to supplementation with thiamin and pantothenic acid derivatives. European Journal of Applied Physiology 77, 486–491 (1998).

73. L. M. Forbes et al., Red Blood Cell Metabolic Responses during Acute Hypoxic Exercise in Healthy Adults. Am J Respir Cell Mol Biol 72, 456–459 (2025).

74. A. A. Barber, F. Bernheim, Lipid peroxidation: its measurement, occurrence, and significance in animal tissues. Adv Gerontol Res 2, 355–403 (1967).

75. W. Kim et al., Polyunsaturated Fatty Acid Desaturation Is a Mechanism for Glycolytic NAD(+) Recycling. Cell Metab 29, 856–870.e857 (2019).

76. R. Carin et al., Effects of Three Different Distance/Elevation Gain Ultra-Trail Races on Red Blood Cell Oxidative Stress and Senescence, and Blood Rheology. Med Sci Sports Exerc 57, 1081–1091 (2025).

77. A. Shenkin, Trace elements and inflammatory response: implications for nutritional support. Nutrition 11, 100–105 (1995).

78. K. F. Adams, G. Johnson, Jr., K. E. Hornowski, T. H. Lineberger, The effect of copper on erythrocyte deformability: a possible mechanism of hemolysis in acute copper intoxication. Biochim Biophys Acta 550, 279–287 (1979).

79. B. O. Adele et al., Alterations in plasma and erythrocyte membrane fatty acid composition following exposure to toxic copper level affect membrane deformability and fluidity in female wistar rats. J Trace Elem Med Biol 80, 127316 (2023).

80. H. Li et al., Mechanics of diseased red blood cells in human spleen and consequences for hereditary blood disorders. Proceedings of the National Academy of Sciences 115, 9574–9579 (2018).

81. R. K. Powers et al., Trisomy 21 activates the kynurenine pathway via increased dosage of interferon receptors. Nat Commun 10, 4766 (2019).

82. H. Zheng et al., C-Reactive protein and the kynurenic acid to quinolinic acid ratio are independently associated with white matter integrity in major depressive disorder. Brain Behav Immun 105, 180–189 (2022).

83. T. Thomas et al., Evidence of Structural Protein Damage and Membrane Lipid Remodeling in Red Blood Cells from COVID-19 Patients. Journal of Proteome Research 19, 4455–4469 (2020).

84. D. A. Bizjak et al., SARS-CoV-2 Altered Hemorheological and Hematological Parameters during One-Month Observation Period in Critically Ill COVID-19 Patients. Int J Mol Sci 23, (2022).

85. E. Nader et al., Increased blood viscosity and red blood cell aggregation in patients with COVID-19. Am J Hematol 97, 283–292 (2022).

86. T. Nemkov, J. A. Reisz, S. Gehrke, K. C. Hansen, A. D’Alessandro, High-Throughput Metabolomics: Isocratic and Gradient Mass Spectrometry-Based Methods. Methods in Molecular Biology (Clifton, N.J.) 1978, 13–26 (2019).

87. J. A. Reisz, C. Zheng, A. D’Alessandro, T. Nemkov, Untargeted and Semi-targeted Lipid Analysis of Biological Samples Using Mass Spectrometry-Based Metabolomics. Methods in Molecular Biology (Clifton, N.J.) 1978, 121–135 (2019).

88. Z. Pang et al., Using MetaboAnalyst 5.0 for LC-HRMS spectra processing, multi-omics integration and covariate adjustment of global metabolomics data. Nat Protoc 17, 1735–1761 (2022).

89. P. Shannon et al., Cytoscape: a software environment for integrated models of biomolecular interaction networks. Genome Res 13, 2498–2504 (2003).

90. D. Stephenson, T. Nemkov, S. M. Qadri, W. P. Sheffield, A. D’Alessandro, Inductively-Coupled Plasma Mass Spectrometry-Novel Insights From an Old Technology Into Stressed Red Blood Cell Physiology. Front Physiol 13, 828087 (2022).

91. S. X. Ge, D. Jung, R. Yao, ShinyGO: a graphical gene-set enrichment tool for animals and plants. Bioinformatics 36, 2628–2629 (2019).

92. W. Luo, C. Brouwer, Pathview: an R/Bioconductor package for pathway-based data integration and visualization. Bioinformatics 29, 1830–1831 (2013).

