## Supplementary Figures for "Long-Distance Trail Running Induces Inflammatory-Associated Protein, Lipid, and Purine Oxidation in Red Blood Cells"

Title

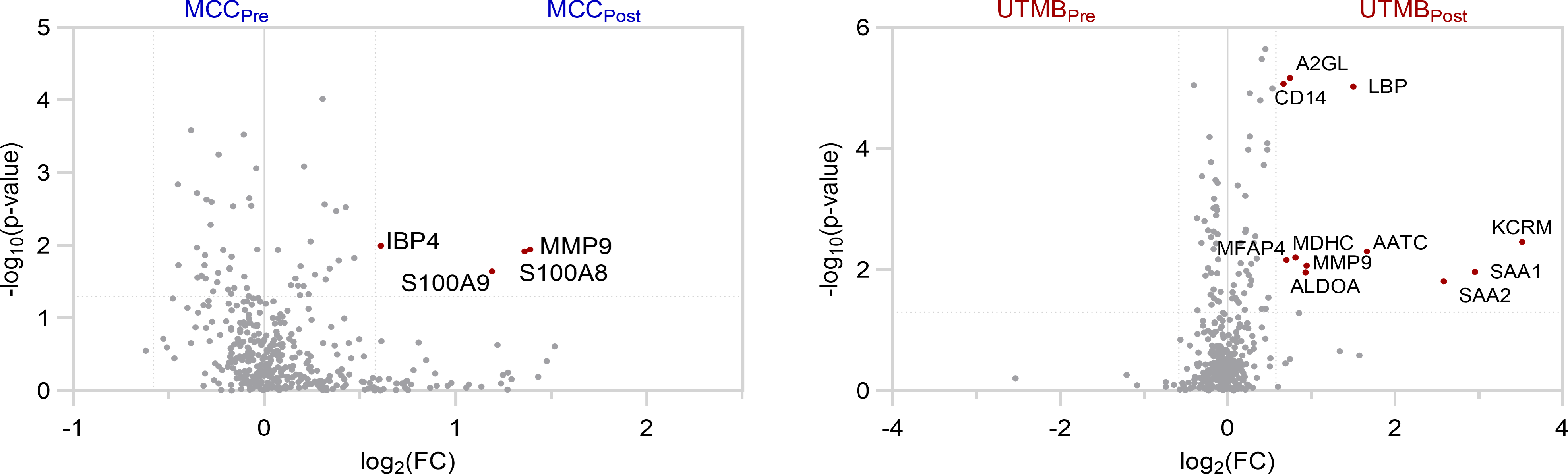

**Supplementary Figure 1 Significant Proteomics Differences Before and After the MCC (left) and UTMB (right)**

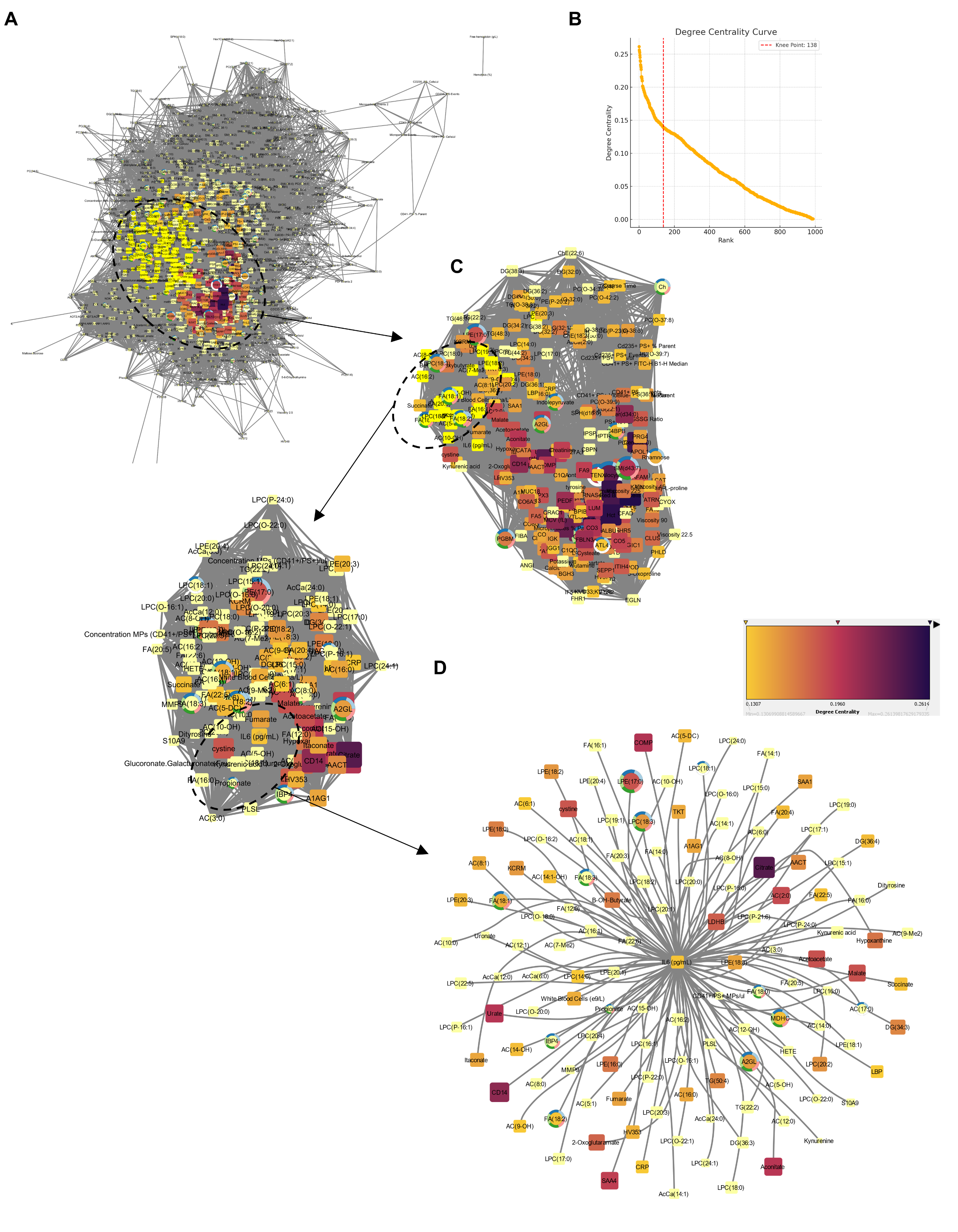

**Supplementary Figure 2 Expanded Debiased Sparse Partial Correlation (DSPC) network for IL-6.** (A) The total network using a degree filter of 1.5, betweenness filter of 1.0, and correlation filter of p < 0.001. (B) Using a Knee Point of 138, the (C) DSPC was refined (C) and centered around IL-6 as the central node (D).

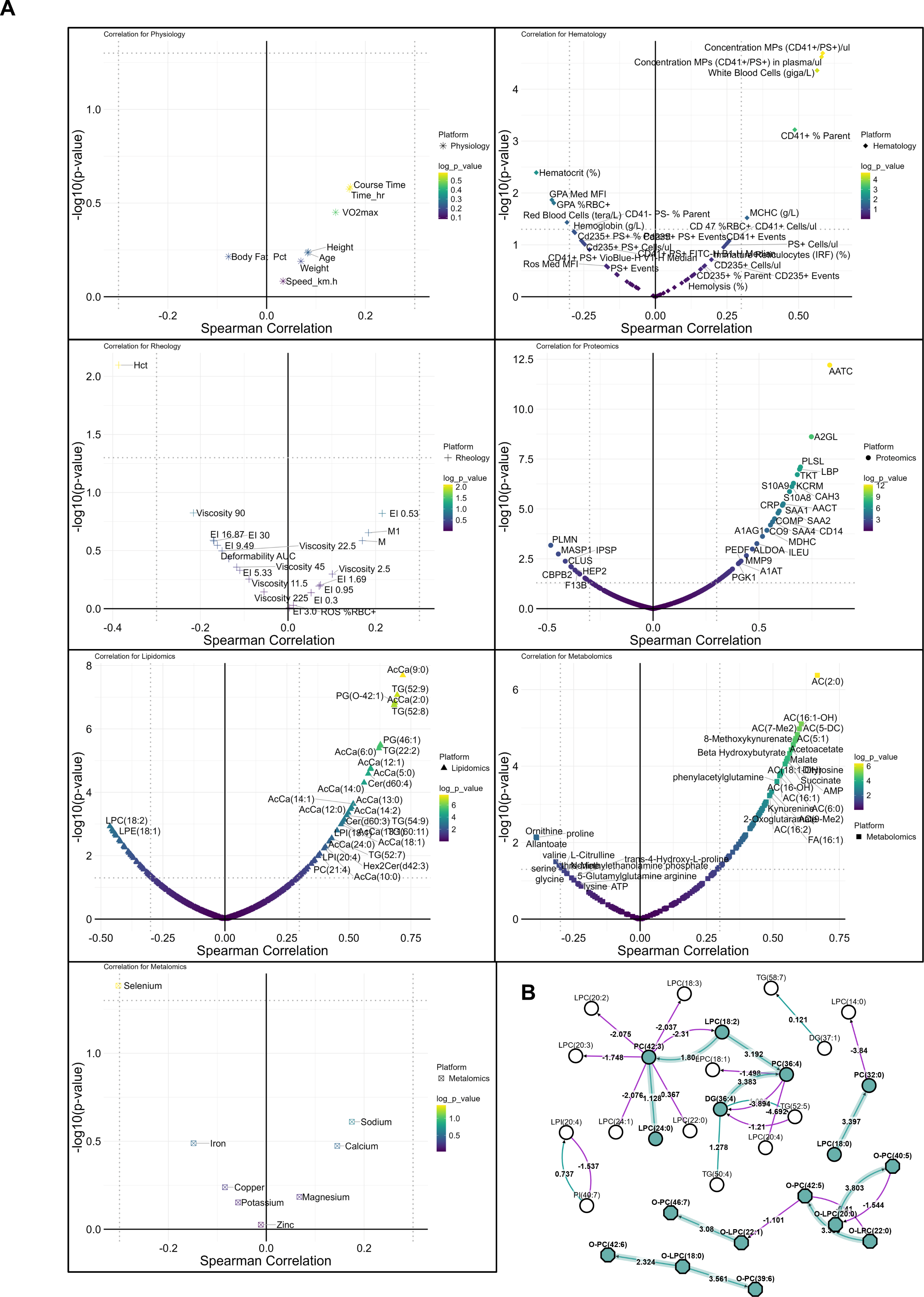

**Supplementary Figure 3 Molecular correlates in plasma with IL-6 as a function of each measurement platform.** (A) Individual correlates with Physiology (*), Hematology (◆), Rheology (+), Proteomics (●), Lipidomics (▲), Metabolomics (■), and ICP-MS Metabolomics (⛝) are shown. (B) BioPAN model of individual lipids is shown.

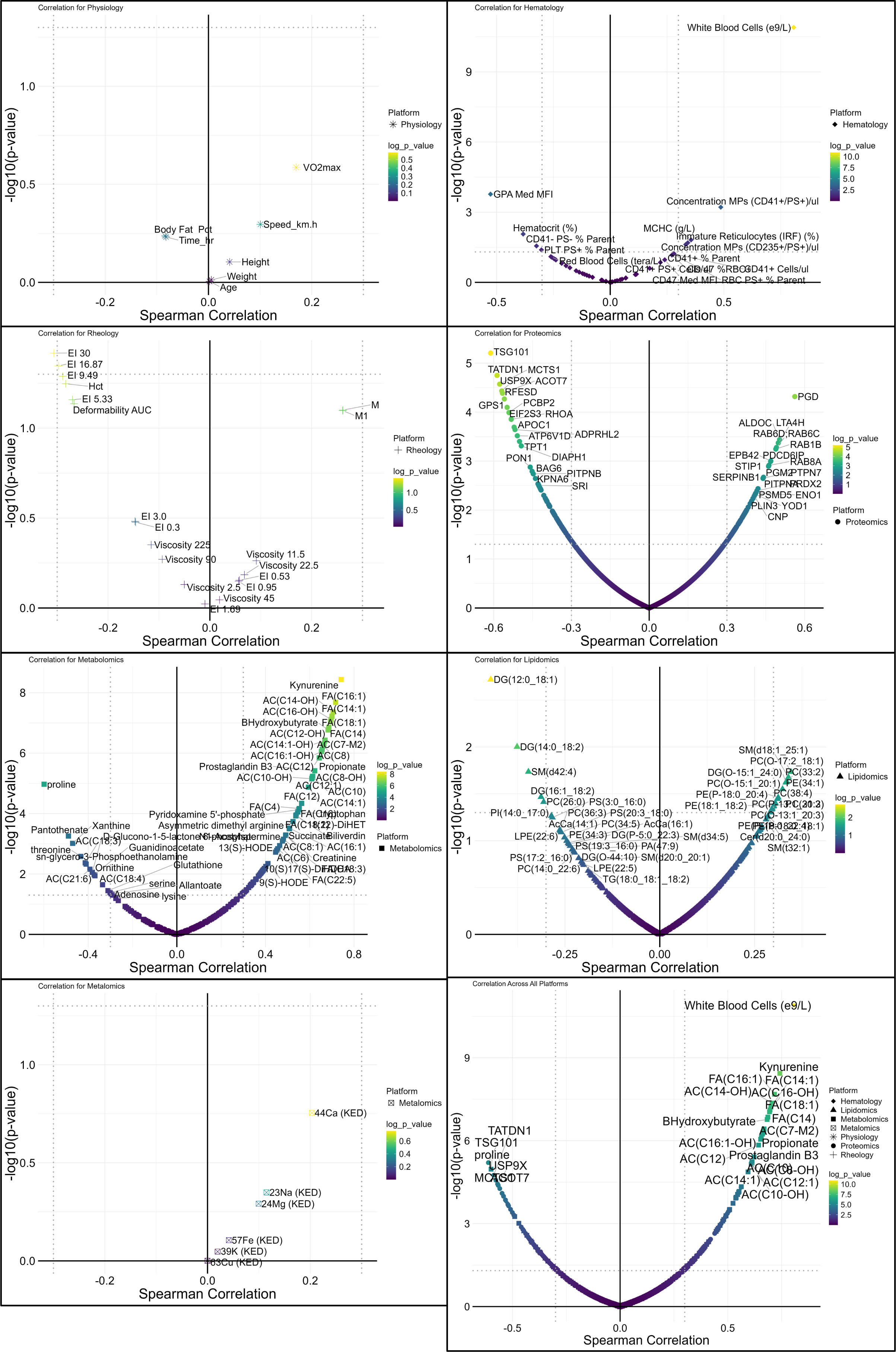

**Supplementary Figure 4 Molecular correlates in RBC with IL-6 as a function of each measurement platform.** (A) Individual correlates with Physiology (*), Hematology (◆), Rheology (+), Proteomics (●), Lipidomics (▲), Metabolomics (■), and ICP-MS Metabolomics (⛝) are shown. (B) BioPAN model of individual lipids is shown.

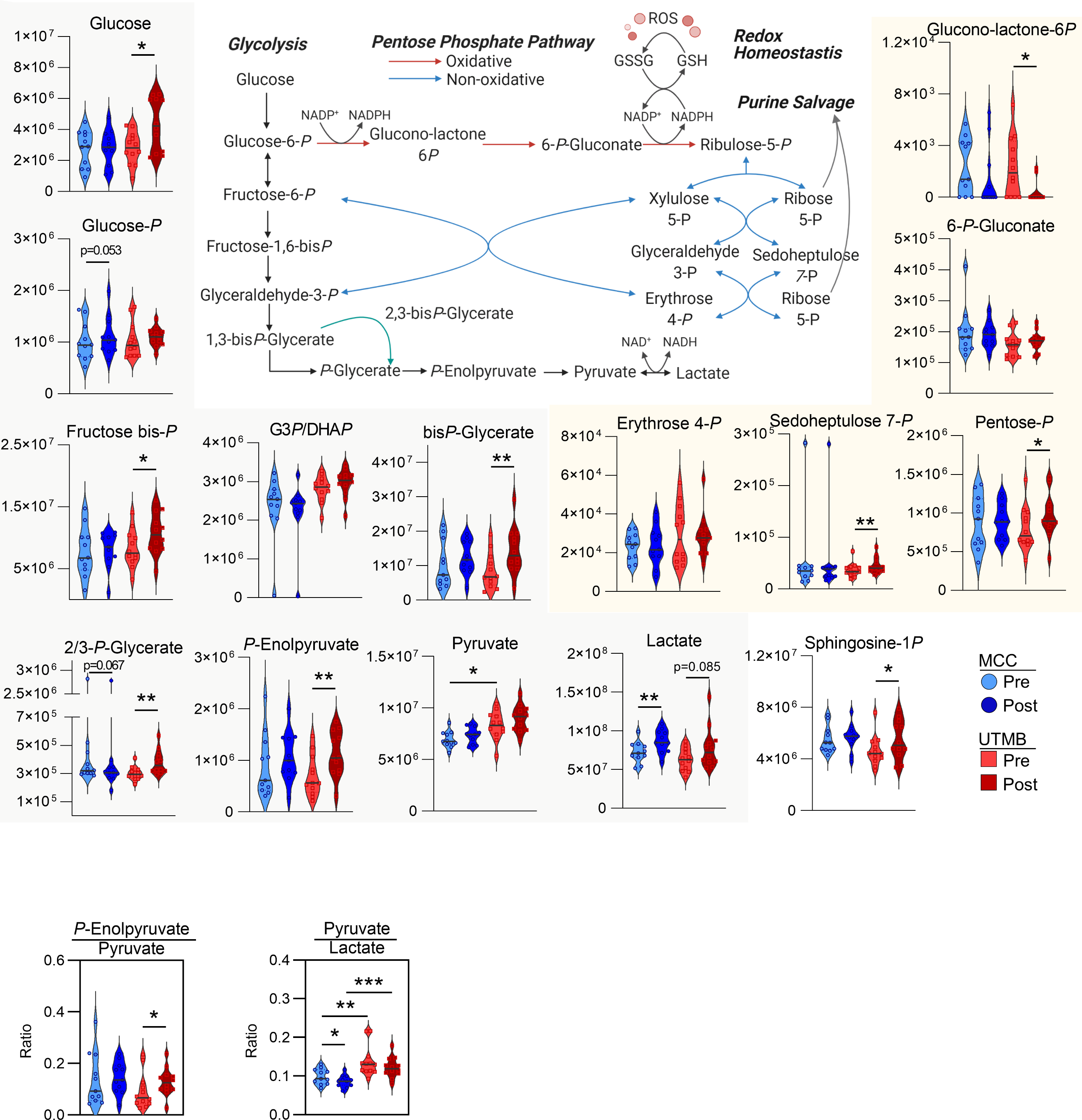

**Supplementary Figure 5 Glycolysis and the Pentose Phosphate Pathway.** Metabolites in glycolysis (gray box) and the Pentose Phosphate Pathway (tan box) are shown. Individual samples from MCC (O) at Pre (light blue) and Post (dark blue), or UTMB () at Pre (light red) or Post (dark red) are shown within violin plots. p-values from Two-tailed homoscedastic T-tests are indicated as *, p<0.05; **, p<0.01, ***, p<0.001; ****, p<0.0001. The pathway map (center) was created with Biorender.com.

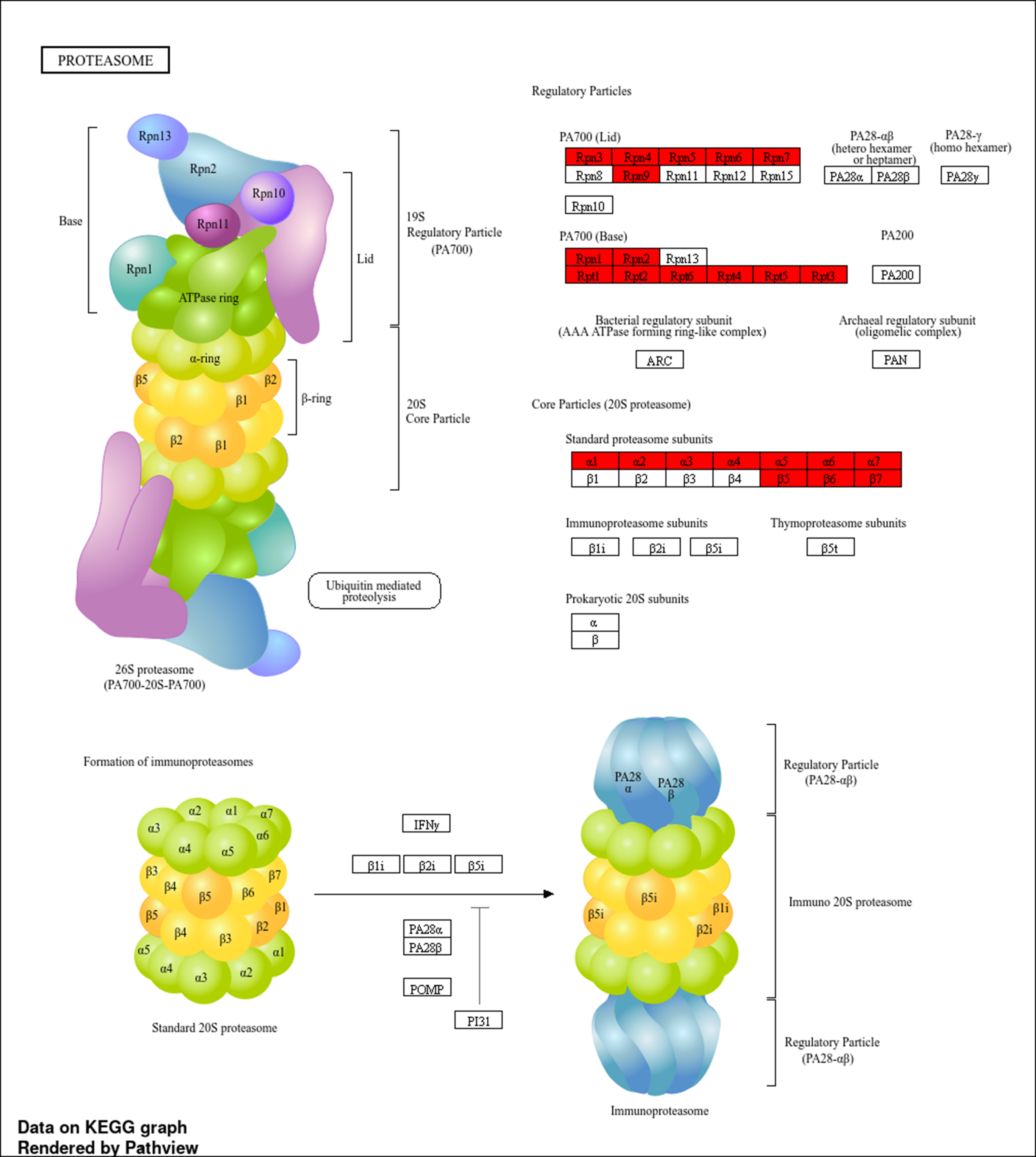

**Supplementary Figure 6 Methionine oxidation of Proteasome regulatory particles.** Proteins with statistically significant change in methionine oxidation (ANOVA Fisher’s LSD FDR < 0.05) are shown in red.

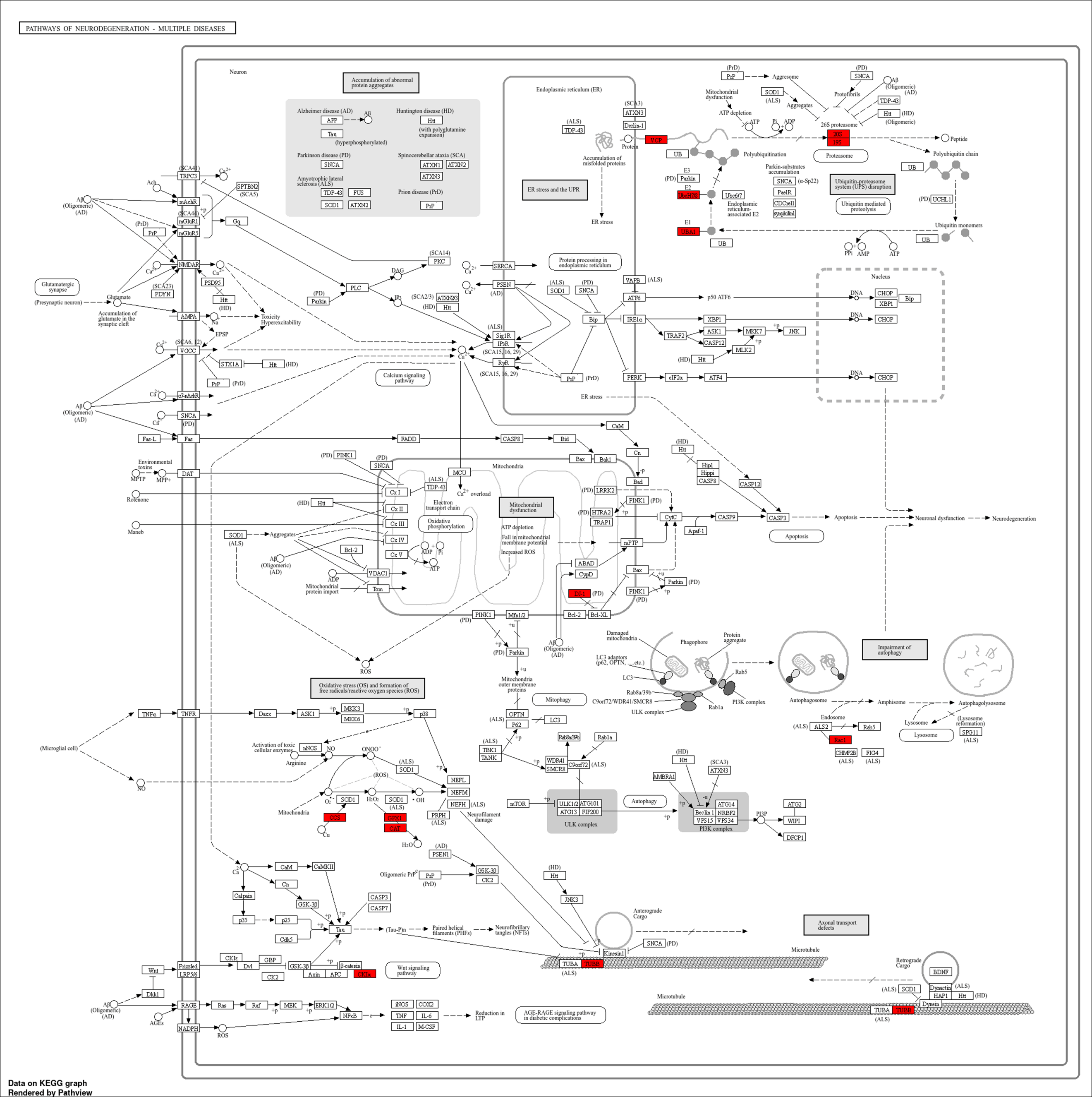

**Supplementary Figure 7 Shared oxidized proteins across multiple neurodegeneration pathways.** Proteins with statistically significant change in methionine oxidation (ANOVA Fisher’s LSD FDR < 0.05) are shown in red.
